# Operationalising LLM-assisted screening of literature to support systematic reviews

**DOI:** 10.64898/2026.07.29.741127

**Authors:** Scott Spillias, Laura Avila-Turriago, Christopher Brown, Ariane Easton, Jack Roberts, Michael Sievers, Stephen Swearer, Andrew Taylor, Brigette Wright, Valeriya Komyakova

## Abstract

Large language models (LLMs) can ease the work of screening titles and abstracts for systematic reviews, but obtaining reliable results requires researchers to make practical choices about which LLMs to use, how to combine their scores into a ranking, and how far down that ranking to read. We aimed to identify a general-purpose workflow that screens accurately, minimises human review effort, and generalises across environmental literature corpora. We ran an ensemble of five open-source LLMs across ten human-annotated systematic reviews from the field of ecology and environmental science spanning 19,155 studies. We then asked: (1) how well an ensemble of LLMs ranks relevant papers above irrelevant ones, and (2) where a human reviewer should stop working down that ranked list. A four-LLM ensemble chosen without any labels came close, on every review, to the best ranking achievable with that review’s annotations (mean Average Precision 0.66 versus 0.68). We tested different rules for when to stop human review, finding that a single *SAFE* stopping rule chosen in advance recovered ≥ 95% of relevant records on all ten reviews while requiring a human to screen 56% of the corpus on average. Against the established open-source active-learning tool `ASReview`, this label-free ranking reached the same recall target at lower human workload on seven of the ten reviews (39% versus 42% of the corpus on average). The paper offers a complete workflow that can be adopted for new, unlabelled reviews, using open-source LLMs small enough to run on a high-end consumer laptop, and we provide it as an open-source R package.

## Introduction

Large Language Models (LLMs) are increasingly being used to support systematic reviews, offering new opportunities to streamline and scale evidence synthesis (Scherbakov et al., 2025; Berger-Tal et al., 2024; Spillias et al., 2024). The ability of LLMs to speed up literature reviews means that LLM-assisted reviews can synthesise a larger number of studies on a shorter timeline than purely human teams. This creates opportunities to more rapidly integrate science into management decisions (Nie and Liu, 2025), to use LLM-powered literature reviews to seed the generation of interpretable mathematical models (Spillias et al., 2026), and to synthesise much larger bodies of literature to identify emergent trends (Chang et al., 2025). One of the most promising applications of LLMs to literature reviews is in the screening phase, where LLMs assist in evaluating large volumes of titles and abstracts to identify their relevance for further detailed review. An effective LLM screening workflow would minimise researcher effort but maintain high recall (i.e., capturing all relevant studies). Initial research has shown that different LLMs display variable competencies in facilitating such screening tasks (Li et al., 2024), in part because it is unclear how to optimise the many workflow choices that balance human screening effort against high recall.

The first choice is what to ask the LLM to output. Existing approaches span a spectrum: at one end, a binary include/exclude decision (Cao et al., 2024; Delgado-Chaves et al., 2025; Oami et al., 2025; Guo et al., 2024; Syriani et al., 2024); in between, a three-way decision that adds a “neutral” option to express uncertainty (Akinseloyin et al., 2026; Thomas et al., 2024; Spillias et al., 2024); and at the other end, a fully continuous relevance score (Dennstädt et al., 2024; Issaiy et al., 2024; Nykvist et al., 2025; Jaumann et al., 2025).

The latter is attractive for human-in-the-loop applications because it preserves the uncertainty that a binary decision discards and because it allows for a meaningful ranked list that a human reviewer can inspect, rather than a binary decision that ignores grey interpretations that are common in literature synthesis screening.

However, a continuous score presents two linked decisions that must be addressed before screening can proceed. The first is how to optimally use LLMs to rank studies, and the second is how far down the LLM-generated ranking to read. The first, which we refer to as the *ranking question*, is how to combine LLM scores into a single ordering of the corpus. The value of multiple screeners is well established in human-only reviews (Stoll et al., 2019; Waffenschmidt et al., 2019), and recent work has extended this to multi-LLM screening strategies (Akinseloyin et al., 2026; Zhang et al., 2025; Sanghera et al., 2025; Thode et al., 2025; Song et al., 2026). These multi-agent methods require aggregation strategies, such as majority voting (Akinseloyin et al., 2026; Zhang et al., 2025), confidence-weighted voting (Song et al., 2026), conservative inclusion (Oami et al., 2025; Thode et al., 2025), or more complex ensemble techniques like random forests (Zhang et al., 2025) and adjudication by a more powerful LLM (Akinseloyin et al., 2026).

The second, the *stopping question*, is how far down the accepted ranking a human reviewer should read. A relevance score must eventually be turned into an include/exclude boundary, yet many studies apply arbitrary thresholds (Dennstädt et al., 2024; Nykvist et al., 2025) or do not specify how thresholds are chosen (Jaumann et al., 2025). More generally than for LLM-assisted reviews, this ranking-and-stopping problem has a long history in information retrieval, where it is studied as “technologically assisted review” (TAR): ranking documents for a reviewer and deciding when to stop reading (Kanoulas et al., 2019; Wang et al., 2022).

Few studies have tackled ranking, stopping, and multi-LLM ensembling simultaneously. The closest is Wang et al. (2024), who combine an ensemble of open-source generative LLMs with a BioBERT classifier and calibrate a score threshold to reach a target recall on biomedical CLEF collections. They test two calibration methods: one extrapolates a median threshold from a set of already-labelled reviews, the other lowers the threshold to the score of known “seed” included studies. Both depend on labelled data that a fresh review may not have, and both produce a fixed inclusion threshold, not a rule for deciding when to stop screening, which is one of the two questions we take on here.

In this study, we develop a protocol for LLM-assisted literature screening that balances human effort against recall. Unlike other multi-LLM screening work (Nawrath et al., 2026), which relies on proprietary frontier LLMs accessed through paid APIs (e.g., *GPT-4* and successors, *Claude*, *Gemini*), our ranker is an ensemble of open-source LLMs that can run locally on consumer hardware. Our test collection is also ecological which has received less attention than other disciplines. We test a range of protocols on ten systematic reviews that received complete human review, from the environmental and ecological domains, together spanning 19,155 records (studies with pre-existing human review decisions). Adopting the rank-then-stop framing from the TAR tradition, we treat the ranking and stopping questions separately. This approach allows us to evaluate a ranking on its own, without first having to pick an include/exclude cutoff, by scoring it with Average Precision (AP), a 0-to-1 summary of how cleanly a ranking places relevant papers above irrelevant ones. We then evaluate a range of stopping rules against the best-performing ranking strategy. The result is an end-to-end, label-free screening workflow with defensible defaults, which we benchmark against the established open-source incumbent for this task, *ASReview* active learning, and release as a reusable R package, *screenllm*.

## Methods

### Overview

We separated LLM-assisted screening into the two questions introduced above, *ranking* and *stopping*, and evaluated each on the same ten labelled reviews (Figure 1). First, we screened every review with an ensemble of five open-source LLMs, each independently scoring every record three times to capture replicate-to-replicate variability. We addressed the *ranking question* (how to order records so the relevant ones come first) by identifying an optimal ranking strategy for each review, then a single “universal” strategy that transferred across all ten reviews, and quantified the performance loss between the universal and review-specific optima. We then addressed the *stopping question* (how far down the ranked list to read) by comparing stopping rules against the universal strategy on each dataset. For both questions we compare against a reference that requires the human labels to compute: for ranking the *best achievable AP*, the highest AP reachable when a review’s human labels are known; for stopping the *ideal stopping point*, the least screening achievable given the true number of relevant records.

**Figure 1:**
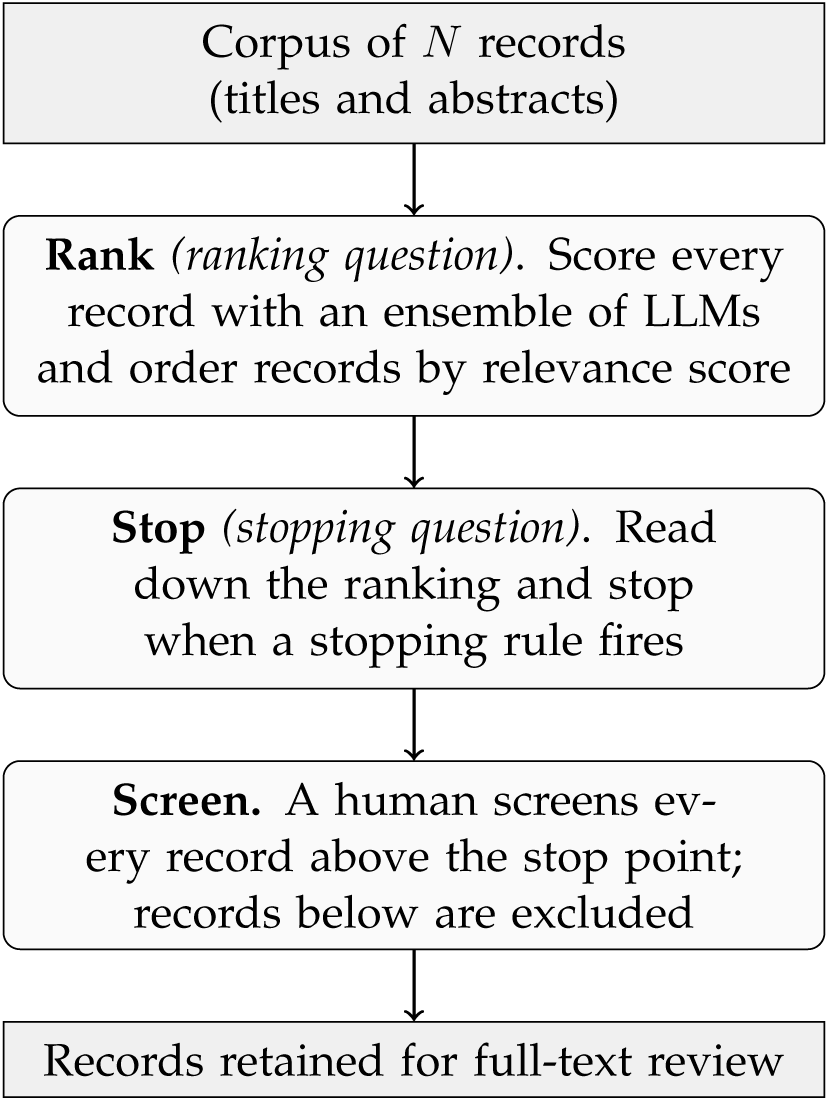
The end-to-end LLM-assisted screening pipeline. The two questions map onto two steps: *ranking* orders the corpus with an LLM ensemble, and *stopping* reads down that ranking until a stopping rule triggers. Throughout, the LLM ensemble triages rather than replaces the human screener: a person still reads every record above the stop point.

### Benchmark reviews

We assembled ten labelled systematic reviews, some in progress and some recently published, from across ecology and environmental science (Table 1). All reviews were screened by trained reviewers using inclusion/exclusion criteria developed to answer a study-specific research question, and each record carries the reviewer’s include/exclude decision. We call those decisions the *human labels*, and use them throughout the paper as the reference against which we measure the LLM ensemble. Review sizes range from 285 to 3,877 human-labelled records, and accept prevalence (the fraction of records that are relevant) varied from 1.9% to 24.5%, providing realistic variation in screening difficulty.

**Table 1:** Labelled systematic reviews used for evaluation. “Records” is the total number of labelled records in the review; “Accepts” is the number of records the human review leads marked relevant; Prev. is the accept prevalence (Accepts / Records); “Rev.” is the number of human screeners who labelled each record (1, 2, or – where unspecified). “Source” cites the published paper for each review where one exists, or marks the review as in preparation. Every label is a screening decision made by human reviewers; for *CBFM*, those decisions came from a human–AI collaborative workflow (Spillias et al., 2024). The full inclusion criteria for each review are reproduced in Supplement P (Per-review inclusion criteria).

| Review | Records | Accepts | Prev. | Rev. | Topic | Source |
| --- | --- | --- | --- | --- | --- | --- |
| Artificial | 3,844 | 73 | 1.9% | 1 | Fitness/morphometric responses to marine habitat type or condition (artificial vs. natural; complex vs. simple) | In prep |
| CBFM | 1,096 | 101 | 9.2% | 5 | Community-based fisheries management in Pacific Island contexts | Spillias et al. (2024) |
| Coastal Wetlands Megafauna | 2,628 | 340 | 12.9% | 1 | Marine megafauna associations with vegetated coastal wetlands | Sievers et al. (2019) |
| Coral Restoration | 3,315 | 340 | 10.3% | 1 | Field-based active coral reef restoration interventions and performance | In prep |
| Habitat Effect | 364 | 52 | 14.3% | 2 | Surrounding-habitat effects on artificial reef performance for fish | In prep |
| Habitat Restoration | 2,827 | 228 | 8.1% | 1 | Organism responses to habitat-forming-species restoration in coastal/marine ecosystems | Sievers et al. (2024) |
| Invasive | 597 | 90 | 15.1% | 2 | Invasive species occupancy of artificial vs. natural marine structures | Avila-Turriago et al. (2026) |
| Light Pollution | 285 | 30 | 10.5% | 1 | Light pollution effects on fitness of marine fish/invertebrates | Easton et al. (2024) |
| Noise Pollution | 322 | 79 | 24.5% | 1 | Marine noise pollution effects on fitness of fish/invertebrates | Easton et al. (2024) |
| Spatial Context | 3,877 | 456 | 11.8% | 1 | Seascape spatial configuration and fitness of marine fish/invertebrates | In prep |

### Out-of-domain reviews

To test whether the workflow generalises beyond marine and coastal ecology, we added two out-of-domain systematic reviews, each the subject of a published LLM-screening study, with the added benefit of comparing our open-source ensemble against proprietary frontier models on the same records and the same human labels. Nawrath et al. (2026) screened a 500-record review of urban greenspace and mental health (8 relevant, 1.6% prevalence) with five proprietary models, and Nykvist et al. (2025) screened a ∼ 12,000-record review of electric-vehicle charging infrastructure (3.7% relevant at the title-and-abstract stage) with *GPT-4*. We ran our recommended “universal” strategy on both corpora using the same prompt template, replicate count, and universal ranking as for the ten marine reviews and scored the resulting rankings with the same metrics (Results, “Generalisation beyond the marine benchmark”; the two external corpora are documented in Supplement O).

### LLM screening prompts and inclusion criteria

The criteria for the accept decision were different for each review. At screening time, each record’s title, abstract, and review-specific inclusion criteria were inserted into a shared prompt template (reproduced in Supplement N) and presented to each LLM. The template asks the LLM to score each criterion separately and then sum those per-criterion scores into a single 0–100 relevance score, where 100 means every criterion is fully met. Each of the review’s *n* criteria carries an equal share of the total (≈ 100*/n* points), and the LLM awards partial credit when a record only partly meets a criterion or the abstract is ambiguous. For habitat-focused reviews (*Artificial*, *Habitat Effect*, *Habitat Restoration*), the prompt also includes a short set of clarifying definitions for what counts as a “habitat”. The same scoring scheme is used for every review, with only the criteria themselves differing (Supplement P).

The LLM screening criteria were derived from those used in each original human-led review, typically written in free-text form and restructured into numbered lists for insertion into the prompt template.

We handled the ten reviews in two modes. For three of them (*Artificial* and *Invasive* during this study, and *CBFM* in Spillias et al. (2024)) we ran a pilot-and-refine loop on the criteria, to show the kind of iteration an attentive review team can do before committing: we screened a small subsample (100 records), examined the LLM’s written justifications, and used *GPT-5* via Microsoft Copilot (August 2025) to fix the recurring failure modes they surfaced (e.g., example lists read as closed, hallucinated exclusion clauses, and undefined domain terms such as “artificial structure” or “natural reef”). The remaining seven reviews were screened with their criteria reformatted but otherwise unchanged, giving an out-of-the-box test on wording that was not tuned to the LLM. We treat the resulting variation as a candidate source of the per-review differences in ranking quality (see Limitations). To quantify how much these choices matter, we separately explored the recommended ensemble’s sensitivity to prompt design (a single holistic score and a binary include/exclude prompt, against our per-criterion score) and to meaning-preserving rewording of the criteria, on three probe reviews spanning the range of ranking difficulty (*Light Pollution*, *Noise Pollution*, and *CBFM*; see Supplement E).

#### Expert re-review and blind re-audit of contested records

While building the benchmark, the ensemble’s confident disagreements with the original *Habitat Effect* screener prompted an expert re-review of the 50 records on which the ensemble and the screener most disagreed. The expert reversed 14 of the original labels in the ensemble’s favour (8 records the screener had accepted in error and 6 it had rejected in error), bringing *Habitat Effect* from 54 to 52 accepts. We adopt these 14 corrected labels in all *Habitat Effect* results reported here.

Because original human labelling is itself a possible source of error driving our results, we also ran a broader blinded re-audit of three reviews (*Light Pollution*, *Spatial Context*, and *Coral Restoration*). Two independent reviewers (SS, VK) re-screened stratified samples seeing only the title, abstract, and inclusion criteria, covering records where the ensemble and human disagreed as well as records accepted by both (Supplement D).

### Large-language model ensemble

We used five open-source LLMs of comparable size: *deepseek-r1:14b*, *gpt-oss:20b*, *qwen3:30b-a3b-instruct-2507*, *mistral-small3.2:24b*, and *gemma3:27b*, of which only *deepseek-r1* is a reasoning LLM (it generates extended internal reasoning before producing an answer). All were served via Ollama (v0.11.3) at low (roughly 4-bit) numerical precision; the exact tags, quantisation formats, and Ollama manifest digests are listed in Supplement M. We deliberately chose LLMs small enough to run on a high-end consumer laptop, but served them on an NVIDIA H100 GPU here to accelerate the large number of replicate screens. Post-training quantisation can shift model outputs (Frantar et al., 2022; Dettmers et al., 2022), an effect that may be larger for reasoning LLMs such as *deepseek-r1*; we discuss the implications in the Limitations. To quantify the ranking’s sensitivity to this choice, we re-served three of the models (*qwen3:30b-a3b-instruct-2507*, *mistral-small3.2:24b*, and *gemma3:27b*) at 8-bit (*Q8_0*) and full 16-bit precision and re-scored three reviews (see Supplement E).

We sampled each non-reasoning LLM at temperature 0.1, leaving the remaining generation parameters at their Ollama defaults (*deepseek-r1* uses its built-in reasoning routine). A low but nonzero temperature avoids the identical scores returned at temperature zero while keeping replicate-to-replicate variance small and guarding against spurious responses (Spillias et al., 2024). The prompt requested a JSON array of records and a 0–100 relevance score per record. Each record was evaluated in a separate prompt (one LLM call per record), and outputs were parsed and any malformed JSON re-requested up to three times before treating the record as “no score returned”.

Each review was screened three times by each LLM, giving 15 full screens per review (5 LLMs × 3 replicates). The replicate count was chosen to give a meaningful per-strategy variance estimate (*r* = 3 leaves at least one within-LLM SD per review for every strategy that uses fewer than three replicates) without being computationally infeasible. Each screen yielded one numerical relevance score per record, plus a justification that can be used for future auditing. We treated these scores as continuous estimates of inclusion likelihood and combined them for each record below. Only 138 of the ≈ 234,700 record-level scores (0.06%) failed to parse as a 0–100 integer, almost all from *deepseek-r1* returning an arithmetic expression or range instead of a single integer, or from JSON parsing failures. When one of an LLM’s replicate scores failed this way, we dropped it rather than counting it as zero, so that LLM’s score for the record was the mean of its remaining valid replicates.

### Reporting metrics

We report a small set of metrics across both questions:

- **Recall** (sensitivity): fraction of relevant records recovered.
- **Miss-rate** (Lost Evidence, 1 − recall): fraction of relevant records a stopping rule leaves unscreened, also known as ‘false negatives’, which is far more damaging to a review than an over-included irrelevant one.
- **Average Precision (AP):** a 0-to-1 summary of how well a ranking puts relevant records above irrelevant ones, equal to the mean precision evaluated at each relevant record as one works down the list (the area under the precision–recall curve). AP = 1 means every relevant record is ranked above every irrelevant one, and AP = prevalence is what random ordering achieves. We used AP because it summarises the entire ranking without requiring an include/exclude cutoff.
- **TNR@95** (true-negative rate at 95% recall): once 95% of the relevant records have been recovered, what fraction of the irrelevant records have been correctly skipped? This is equivalent to normalised Work Saved over Sampling (WSS) (Kusa et al., 2023). We used TNR@95 rather than the original WSS@95 (Cohen et al., 2006) because it is bounded in [0, 1] and is comparable across reviews.
- **Workload:** fraction of the corpus screened to reach a given recall target.

Figure 2 shows how these metrics connect the two questions: AP measures how far a ranking front-loads the relevant records, and the workload to reach a recall target follows directly from it.

**Figure 2:**
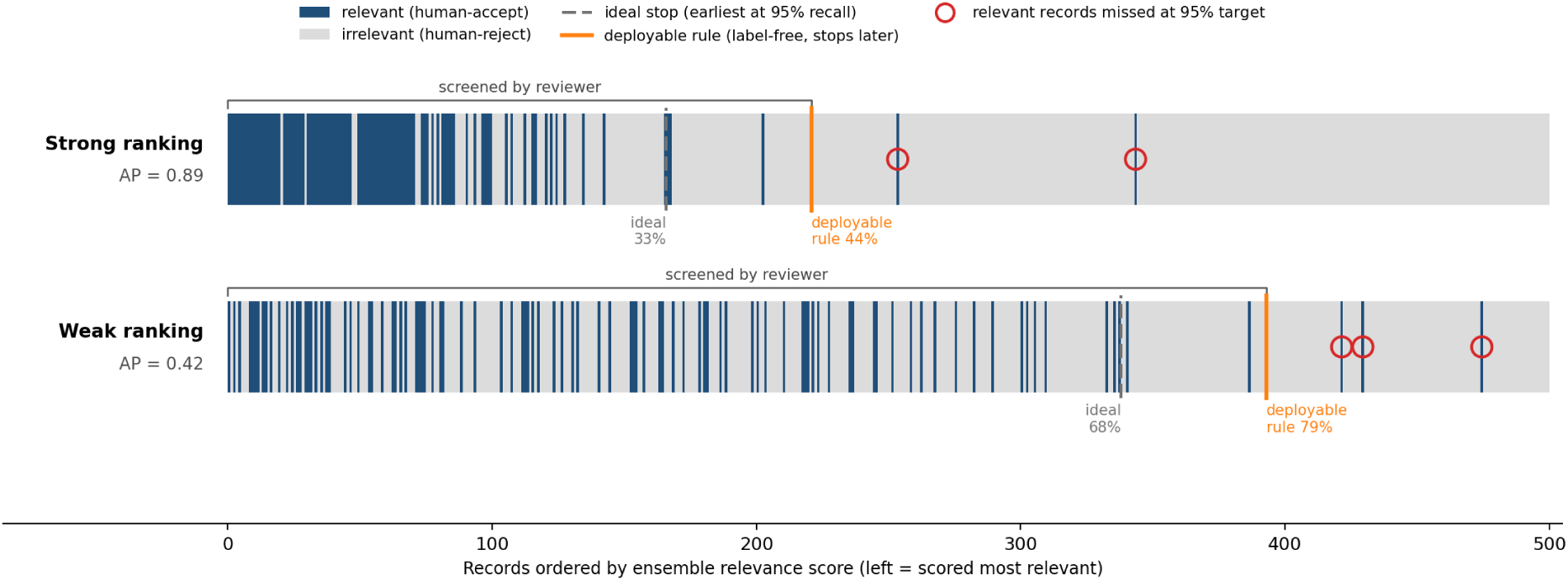
How ranking quality and the stopping decision interact. Each cell is one record, ordered left to right by ensemble relevance score; blue cells are relevant (human-accept), grey cells irrelevant. A strong ranking (top) front-loads the relevant records and so scores a high AP (0.89), whereas a weak ranking (bottom) spreads them and scores low (0.42). Two stopping points are marked on each row: the *ideal* stop (grey dashed), the earliest point that recovers 95% of the relevant records when the labels are known, and a *stopping* rule (orange), which cannot see the labels and so stops somewhat later. A human reviewer screens everything to the left of the deployable stop (the workload); the few relevant records stranded beyond it (circled) are the small fraction the 95% target permits to be missed.

### The ranking question

The ranking question asks how to order the corpus so that relevant records tend to appear earlier. A ranking strategy combines the ensemble’s per-record, per-LLM relevance scores into a single aggregated score *A* per record, which then gives an ordering of the corpus. For aggregating LLM scores, we tried every combination across three dimensions:

- **Which LLMs to include.** All combinations of one or more of the five LLMs (31 in total, 2^5^ − 1).
- **How many of each LLM’s three replicates to use (***r*). Each LLM can contribute one, two, or all three of its replicates (*r* ∈ {1, 2, 3}); when it contributes fewer than three, several replicate combinations are possible (a four-LLM ensemble at one replicate each has 3^4^ = 81), whereas *r* = 3 leaves a single fixed combination.
- **How to combine the scores.** Four ways of pooling the selected scores into one number per record: take the maximum, the arithmetic mean, the median, or the mean of the top two scores (*top-k mean* with *k* = 2).

This gave 31 × 3 × 4 = 372 ranking strategies per review.

For each (review, strategy, replicate combination) we pooled the selected scores into a single aggregated score *A* per record using the strategy’s chosen method. Where we report a per-record standard deviation (*SD*), as a measure of how much the LLMs disagreed on that record, it is taken over exactly the scores that went into *A* (not over all three replicates of every LLM). Records are then ranked from highest *A* to lowest, and we summarise the quality of that ranking with Average Precision (AP).

When a strategy uses fewer than three replicates per LLM (*r <* 3), it admits several replicate combinations. We treated each as one realisation of the strategy and reported the mean AP across them which is what a reviewer would expect, on average, from running the strategy once.

#### A per-review best achievable AP

For each of the ten reviews, we independently identified the ranking strategy that maximised mean AP on that review’s labels. When two strategies were within 0.005 AP, we broke ties first on lower replicate-to-replicate variability and then on the smaller ensemble. This resulting per-review best is the best achievable AP, requiring the labels we are trying to predict. It cannot therefore be used as a recipe in real practice, but instead indicates the ranking quality the ensemble could reach if it were perfectly tuned to that review.

#### A single universal strategy

To approximate what a real reviewer could actually use, we picked a single “universal” ranking strategy, one chosen without seeing any individual review’s labels and judged only on the ranking it produced. For each (review, ranking strategy) pair we used the mean-across-replicates AP from the best-achievable-AP analysis above, averaged AP across reviews, and picked the strategy with the highest cross-review mean as the universal best. This gives a review-agnostic strategy, but with a potential cost that can be calculated by comparing performance against the per-review best AP calculated earlier.

However, because the same ten reviews are used both to pick the universal strategy and to evaluate it, a strategy that happens to fit our particular reviews well could look better than one that would generalise to a new review. To test whether this mattered, we ran a leave-one-out (LOO) cross-validation: we took one of the ten reviews out, picked the universal best from the remaining nine, then evaluated that strategy on the review we had held out, repeating with each review taken out in turn. If different hold-out splits selected different strategies, we could not be confident the strategy is truly universal, but if they all selected the same one, it is more likely to generalise to a new review.

#### How much does the ensemble size matter?

Running extra LLMs costs additional resources, so we looked at how ranking quality changed as we varied the ensemble size from a single LLM up to all five, asking whether the ensemble could be made smaller without giving up much AP and whether performance was still climbing at five. For each ensemble size *s* ∈ {1, 2, 3, 4, 5}, we identified the size-*s* universal best (the size-*s* subset with the highest mean AP across the ten reviews) and evaluated it on each review. We also computed, for each review, the best AP achievable at each size (a within-size best) and tested whether allowing a larger ensemble can ever lower the best AP achievable on that review. Finally, we ran another leave-one-out analysis on the full size-five ensemble to identify the LLM contributing least. We applied the same returns analysis to replicate count, comparing the recommended ensemble’s ranking, screening-to-target workload, and recall at one, two, and three replicates per LLM.

### The stopping question

#### Stopping rules

Using the universal strategy to rank each of the ten reviews, we evaluated seven stopping rules against a target recall of 95% and compared them to an idealised benchmark we call the *ideal stopping point*: a rule that is told in advance exactly how many relevant records the corpus contains, and stops as soon as it has found 95% of them (Callaghan and Müller-Hansen, 2020). Like the best achievable AP for ranking, the ideal stopping point marks the least screening any rule could possibly require. Every rule we tested acts on rank position (i.e., how far down the ordered list a record sits), not on a fixed score threshold, because no single threshold transfers across reviews (Supplement K).

The seven rules fall into three families. **Sequence-only** rules use only the running stream of accept/reject decisions: *consecutive negatives* stops after a long run of irrelevant records, and *percentage* stops after a fixed fraction of the corpus has been read. **Periodic-audit** rules check a small random sample of the unscreened tail every 50 records and update an estimate of how many relevant records remain. We evaluated four such rules. Two are our own, a Bayesian rule and an order-statistics rule formulated from standard sampling statistics; the other two are the frequentist recall-confidence rule of Callaghan and Müller-Hansen (2020) and a Chao1-based variant adapted from Chao (1984). The **combined heuristic** *SAFE* (Boetje and van de Schoot, 2024) stops only when a coverage threshold, a run of consecutive negatives, and a pre-screen estimate of the positive count all agree. Per-rule descriptions and the parameter grids we swept are in Table S11, and formal definitions are in Supplement L. We also tested a fourth family of “model-aware” rules that use the LLM’s relevance score for every record to judge when the target is reached; none improved on SAFE, so we report them only in Supplement Q.

The exact stopping conditions for the audit-based rules use a few standard quantities, including the position in the ranked list, the number of relevant records found so far, and a “recall budget” that captures how many relevant records the unscreened tail can still hold without violating the 95% target. These formal definitions are given in Supplement L.

##### Choosing rule settings

Three of the seven rules require parameters chosen in advance: the run length for consecutive negatives, the fraction for percentage, and both the minimum coverage and the consecutive-negative run length for SAFE (Table S11). For each review and each such rule, we took the lowest-workload setting that still achieves the 95% recall target on that review. Whilst this used the labels in hindsight (a real reviewer would have to commit to a setting in advance), it is an optimistic treatment of each rule and lets the comparison reflect the rule family.

For each review-rule pair we used the universal-strategy ranking and applied the rule’s stopping criterion record by record. The position where it first triggered the rule gave the realised recall, the workload (the fraction of the corpus read before stopping, stop position*/N*), and whether the rule met the 95% target. The audit-based rules (Bayesian, recall-confidence, order-statistics, Chao *r_r_*) have no parameters to tune, so their stopping point was whatever the rule itself decided on each review.

SAFE carries an extra source of variability, because one of its three conditions compares the positives found so far against an estimate from a 200-record spot-check of the corpus, so its stopping point depends on which records happen to fall in that sample. We therefore re-evaluated SAFE the way a reviewer would actually use it, fixing its settings in advance and sweeping a grid of minimum-coverage and run-length values across 500 independent spot-check draws per review (Table S8; Supplement H). The two evaluations judge success differently: in Table S7 a review counts as a success if *any* setting reaches 95% recall, while Table S8 applies the stricter test that at least 95% of the spot-check draws reach recall.

### Comparison with active learning

We benchmarked our ensemble ranking against `ASReview` (van de Schoot et al., 2021), the established active-learning tool for study screening, in which the reviewer marks an initial pilot and the tool re-orders the unscreened records after every decision. We ran it in its default configuration, averaged over ten random pilots, and scored it with the same recall-at-workload measure used for our own ranking(Supplement J).

## Results

### The ranking question

#### A per-review best achievable AP

Across the ten reviews, the best achievable AP varied widely, from 0.47 on *Artificial* to 0.98 on *Noise Pollution* (mean 0.68, median 0.67, 90% confidence interval over reviews [0.61, 0.75]; Table S2).

We found that refining a review’s criteria against a pilot did not translate into a higher best achievable AP, as the three reviews whose criteria we pilot-refined (*Artificial*, *Invasive*, *CBFM*) reached a mean best-achievable AP of 0.63, which was comparable to the 0.70 mean of the seven screened with criteria as supplied, and their variation (0.47–0.76) falls within that of the out-of-the-box reviews (0.54–0.98).

Three patterns recurred across nearly every review (the best ensemble for each review is listed in Table S2 in Supplement C). First, the *mean* aggregation was the best aggregator on 9*/*10 reviews, with *Habitat Effect* the sole exception (it preferred the *median*). *Max* was never best on any review. Second, using all three replicates per LLM was best on 9*/*10 reviews. The sole exception (*Spatial Context*) preferred two replicates, but the gap between two and three was negligible (SD ≤ 0.009 AP). Third, smaller ensembles consistently beat the full five-LLM ensemble. On every review the best size-five ensemble was strictly worse than the best ensemble at some smaller size (gaps from −0.003 on *Coral Restoration* to −0.081 on *Light Pollution*), so no review’s overall best used all five LLMs. Five reviews preferred two-LLM subsets, three preferred three-LLM subsets, and two preferred four-LLM subsets (Supplement C). We return below (ensemble size) to whether this pattern persists in the cross-review analysis.

A high best achievable AP does not automatically mean less screening work for the reviewer. *Artificial* illustrates this as its best achievable AP is the lowest of the ten (0.47), yet its ideal-stop workload-to-95%-recall is also the lowest in the collection (16%) because the very low accept prevalence (1.9%) means the small number of positives can be reached without screening deep into the ranking list. *Habitat Restoration* shows the reverse, with a strong best achievable AP of 0.69 yet one of the highest ideal-stop workloads in the collection (50%), because its relevant records are thinly spread and the last few sit deep in the ranking. These metrics diverge because AP summarises the whole ranking and is dominated by the bulk of relevant records, while workload-to-recall is dominated by the tail of late-ranked positives.

Whether prevalence or review size predict this best-achievable-AP spread is examined in Supplement B.

#### A single universal strategy

We found that the “universal” strategy that maximises mean AP across the ten reviews was the four-LLM *mean* ensemble {*gemma3*, *gpt-oss*, *mistral-small3*, *qwen3*} with all three replicates per LLM. It reached mean AP 0.657 across the ten reviews (90% confidence interval over reviews [0.593, 0.727], SD across reviews 0.137). All four of the runner-up strategies used the same four-LLM subset with two replicates per LLM, swapped one LLM in, or swapped one LLM out, and all sat within 0.010 AP of the winner. The five-LLM ensemble that adds *deepseek-r1* at two replicates per LLM performed just below, with a mean AP 0.651, which is within the replicate noise of the four-LLM best performer.

The cost of using this universal strategy in place of the per-review best achievable AP was small (Table S2, ΔAP column), with the mean per-review gap being 0.022 AP. The worst case was *Light Pollution* at 0.060 AP, while on *Spatial Context* the universal was essentially indistinguishable from the review-specific best achievable AP (Δ = 0.003). This choice is also robust to which reviews we happen to have: in leave-one-out cross-validation (best strategy chosen on nine reviews, tested on the tenth) the same four-LLM *mean* ensemble won all ten tests, at a held-out cost close to the in-sample value (0.020 AP on average; 90% bootstrap interval [0.012, 0.029]). These estimates are also stable to replicate noise and to label error, which we quantify below (“Robustness to analytic choices and label error”).

The strategy is also insensitive to which reviews are published (six of the ten are already published, four in preparation; Table 1). The recommended four-LLM *mean* ensemble was the top-ranked of the 31 candidate subsets on all ten reviews, on the six published alone, and on the four unpublished alone. The LLMs therefore gain no unfair advantage from possibly having the already-published reviews in their training data.

Across all 372 candidate strategies the universal ensemble has the highest mean AP and lies on the *Pareto frontier* (no strategy beats it on both mean AP and worst-review AP; Figure 3), and on a review-by-review basis it sits just below each review’s with-labels best achievable AP.

**Figure 3:**
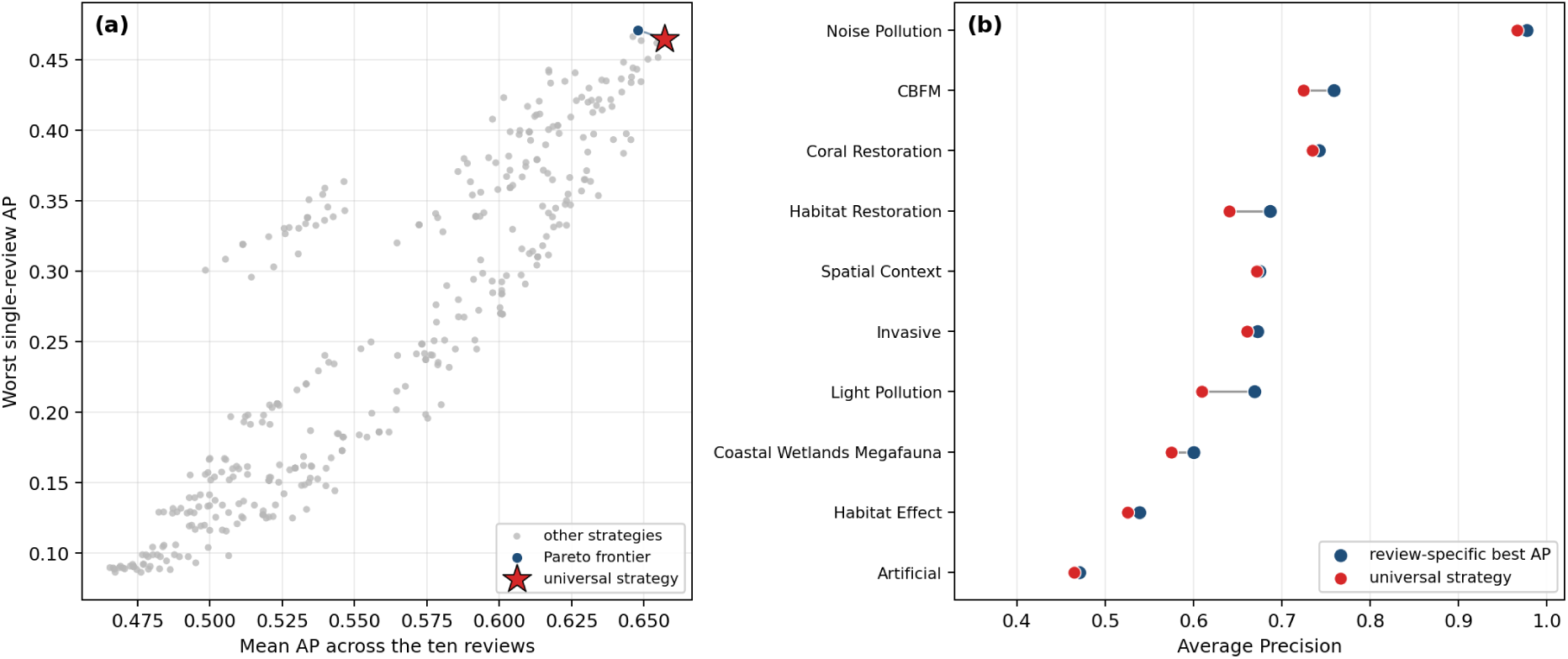
Generality of the universal ranking strategy. **(a)** Each point is one of the 372 candidate strategies, placed by its mean AP across the ten reviews and its worst single-review AP. The universal four-LLM *mean* ensemble (star) has the highest mean AP and lies on the Pareto frontier (blue), so no strategy beats it on both axes. **(b)** For each review (sorted by best achievable AP), the universal strategy’s AP (red) beside that review’s own with-labels best achievable AP (blue). The universal strategy falls just short of the bespoke optimum on every review, with a mean gap of 0.022 AP and a worst gap of 0.060 on *Light Pollution*. Replicate-to-replicate variation in the best achievable AP is negligible on every review, so error bars are not shown.

#### How much does the ensemble size matter?

Returns diminished quickly as the ensemble grew, though any ensemble clearly outperformed a single LLM. The best single LLM (*gpt-oss*) reached mean AP 0.550. Adding *gemma3* to form the best pair raised this by +0.053, adding *mistral-small3* for the best trio a further +0.033, and adding *qwen3* for the best four-LLM set (the universal subset) only +0.006 more, reaching 0.642. A fifth LLM (*deepseek-r1*) subtracted 0.003, which was within replicate-to-replicate noise. The bootstrap 90% interval on the four-versus-five difference [−0.003, +0.010] includes zero, and across the ten reviews individually the universal four-LLM subset outperformed the universal five-LLM subset on six and lost on four. The gain had shrunk to within noise by the fourth LLM and dipped at the fifth, so the ensemble appears to have about reached its ceiling; a sixth similar LLM would be unlikely to help, though we did not test this directly. Replicate count showed the same pattern, as for the recommended four-LLM ensemble, increasing from one replicate per LLM to three raised mean AP by only 0.008 (0.649 to 0.657), which is inside the replicate-to-replicate noise itself, and left both the workload to reach 95% recall and the recall at the advance-fixed SAFE workload essentially unchanged (Supplement F). The ranking benefit therefore comes from ensembling different LLMs, not from repeated sampling of the same ones.

This also held within reviews on average (Figure S4 in Supplement F). On every review, the best achievable AP flattened or dipped slightly once the ensemble passed three or four LLMs, so a larger ensemble was never reliably a better one. From a practical perspective, this means that ensemble size matters more than its exact composition of LLMs, and so a reviewer can use any three or four comparably-sized open-source LLMs they have and can expect most of the benefit of ensembling, without needing to reproduce the specific set of LLMs we report here. Again, the same holds for replicates (Figure S3), but we note that the three replicates we ran are useful for estimating uncertainty and for guarding against an occasional failed run, so a reviewer running a single screen may ultimately decide to use fewer replicates.

#### Robustness to analytic choices and label error

Prompt design mattered in some cases, and our original design was best suited to the task (Supplement E). Collapsing the per-criterion partial-credit prompt to a binary include/exclude decision lowered AP on all three reviews (by 0.02 to 0.04, every 90% interval excluding zero), while a single holistic 0–100 score helped the hardest review (*Light Pollution*, +0.12) but was neutral on the other two. Rewording the inclusion criteria changed AP negligibly (|ΔAP| ≤ 0.018), and serving the three quantisable models at 8-bit or full 16-bit precision instead of 4-bit left the ranking essentially unchanged (mean |ΔAP| versus 4-bit of 0.006 at 8-bit and 0.008 at full precision). The one consequential choice, keeping the per-criterion partial-credit prompt rather than a binary decision, confirms the recommended design.

Beyond these design choices, the reported metrics depend on two quantities we do not observe exactly: the stochastic LLM scores and the human labels. Resampling the four universal models’ replicates changes the estimated universal AP by only about 0.01 (cross-review mean interval [0.651, 0.658]), showing that uncertainty from LLM stochasticity is small relative to the observed performance differences (Supplement D).

Human label error is the second source. An expert re-review of *Habitat Effect* corrected 14 of the 50 records on which the ensemble and the human most disagreed (near 28%), and a stratified blind re-audit of three further reviews (*Light Pollution*, *Spatial Context*, and *Coral Restoration*), each re-screened by two independent reviewers, measured error across the whole score range (Supplement D). On the critical false negatives, those the ensemble scored below 30 against a human Accept, all 11 examined were judged irrelevant by both readers, i.e. over-inclusions by the original screener. On confidently-accepted records, the reviewers disagreed with the original label about half the time, and the two readers disagreed with each other over the genuinely contested records (one upheld the ensemble on about a third, the other on almost none). Propagating the audit’s measured, stratum-specific correction rates through the ranking actually increases the cross-review mean AP slightly to be 0.66–0.69 against the 0.657 original estimate, because the contested corrections favour the ensemble.

#### Generalisation beyond the marine benchmark

The universal ranking transferred to both out-of-domain reviews. On the urban-greenspace review (Nawrath et al., 2026), the ensemble ranked all eight relevant records within the top 18% of the corpus, and on the larger charging-infrastructure review (Nykvist et al., 2025), it ranked such that a reviewer would reach 95% recall after screening 28% of the corpus. Both workloads sit within the 14–58% ideal-stop range spanned by the ten marine reviews.

#### Comparison with active learning

To reach 95% recall, we found that a reviewer using our method would screen 39% of the corpus on average, whereas we found that with ASReview’s active learning method, a reviewer would screen 42%. For 7 of the 10 reviews, our method is the lower-workload option and uses no labelled examples (Supplement J), but does require well-formed screening criteria.

##### Where does the universal ranker misplace relevant records?

For screening, the error that matters most is a false negative, where a genuinely relevant record is rated so poorly by the LLM screen that a human is unlikely to see it. We found that the universal ensemble rarely made this error. Across the ten reviews, only 0.9% of human-accepted records received an aggregated score below 30 on the 0–100 scale under the universal ranking, and on every review confidently-rejected relevant records were rare (per-review counts in Table S1). The blind re-audit (above, “Robustness to analytic choices and label error”) indicates these are rarely egregious misses: the below-30 records it examined were over-inclusions by the original screener, not relevant papers the ensemble lost. Instead, mistakes were overwhelmingly false positives (Figure 4), which is desirable from a human-AI collaboration perspective anyway (Spillias et al., 2024).

**Figure 4:**
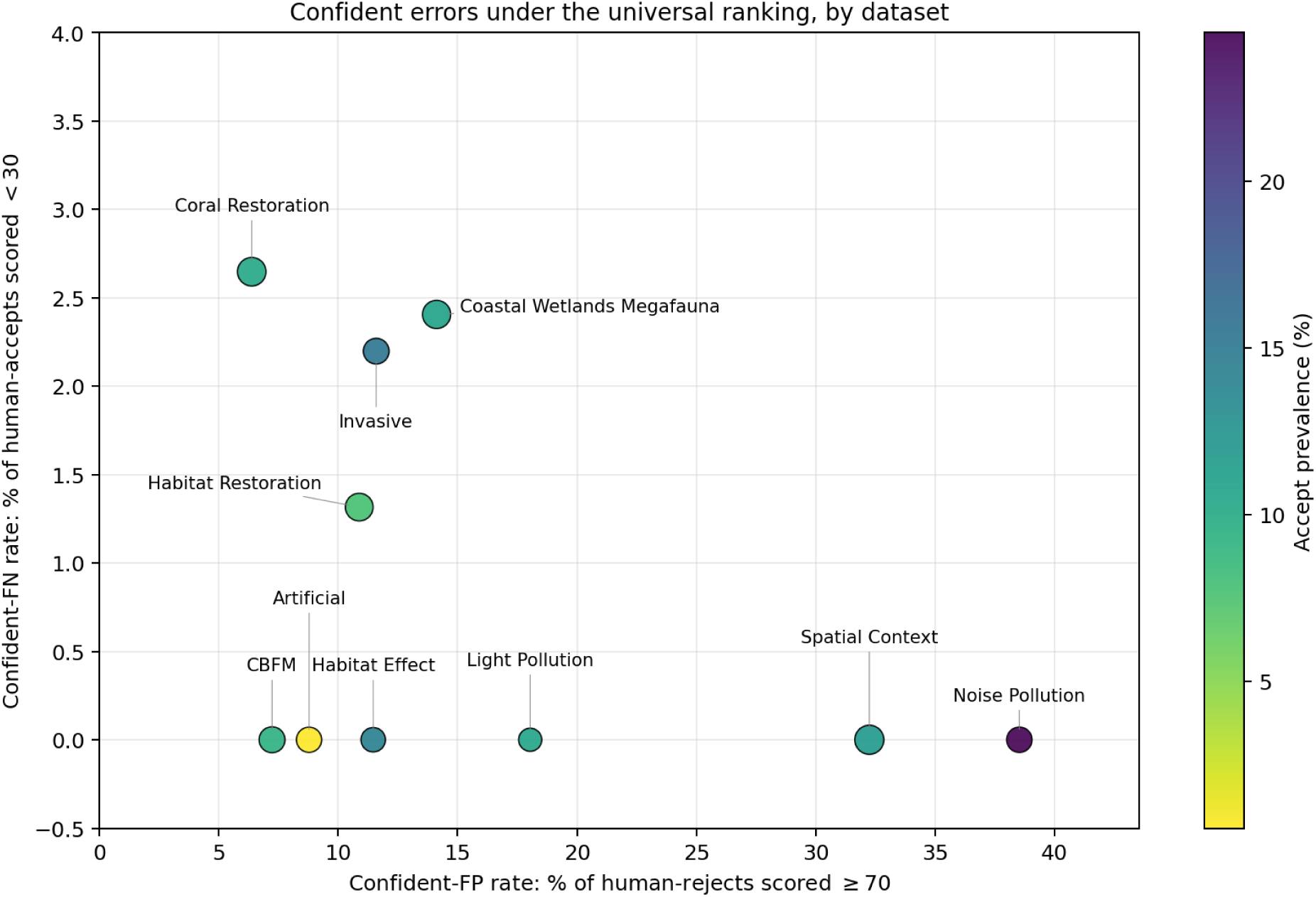
Confident errors of the universal ranking, per review. Each point is one review; marker area scales with the number of human-accepted records and colour with accept prevalence. The horizontal axis is the rate at which the ensemble confidently *accepts* a record humans rejected (LLM score ≥ 70); the vertical axis is the rate at which it confidently *rejects* a record humans accepted (LLM score *<* 30). For a screening workflow the desirable axis is the vertical one: every review sits at or near zero on the confident-FN axis (cross-review mean 0.9%). Confident-FP rates are higher and more variable across reviews, but the asymmetry (the ensemble is far more likely to over-include than to confidently mis-reject) is the relevant property when a human screens everything above the stopping point.

### The stopping question

#### When can a reviewer stop?

We compared the seven candidate rules against the ideal stopping point introduced above, which is the idealised rule that knows the true number of relevant records and stops as soon as it has found 95% of them. Because a real reviewer does not know that number in advance, the ideal stopping point represents the least screening any rule could feasibly require on the universal ranking, and the bar against which we judge the seven candidates.

##### The lower limit on workload

When the true number of relevant records is known, the ideal stopping point reached the 95% recall target on all ten reviews at a mean workload of 39% (median 39%; range 16% on *Artificial* to 58% on *Habitat Effect*; Figure 5). No review required near-complete screening at the ideal stopping point, as all ten reached the target at or below 58% of the corpus, and half at or below 40%.

**Figure 5:**
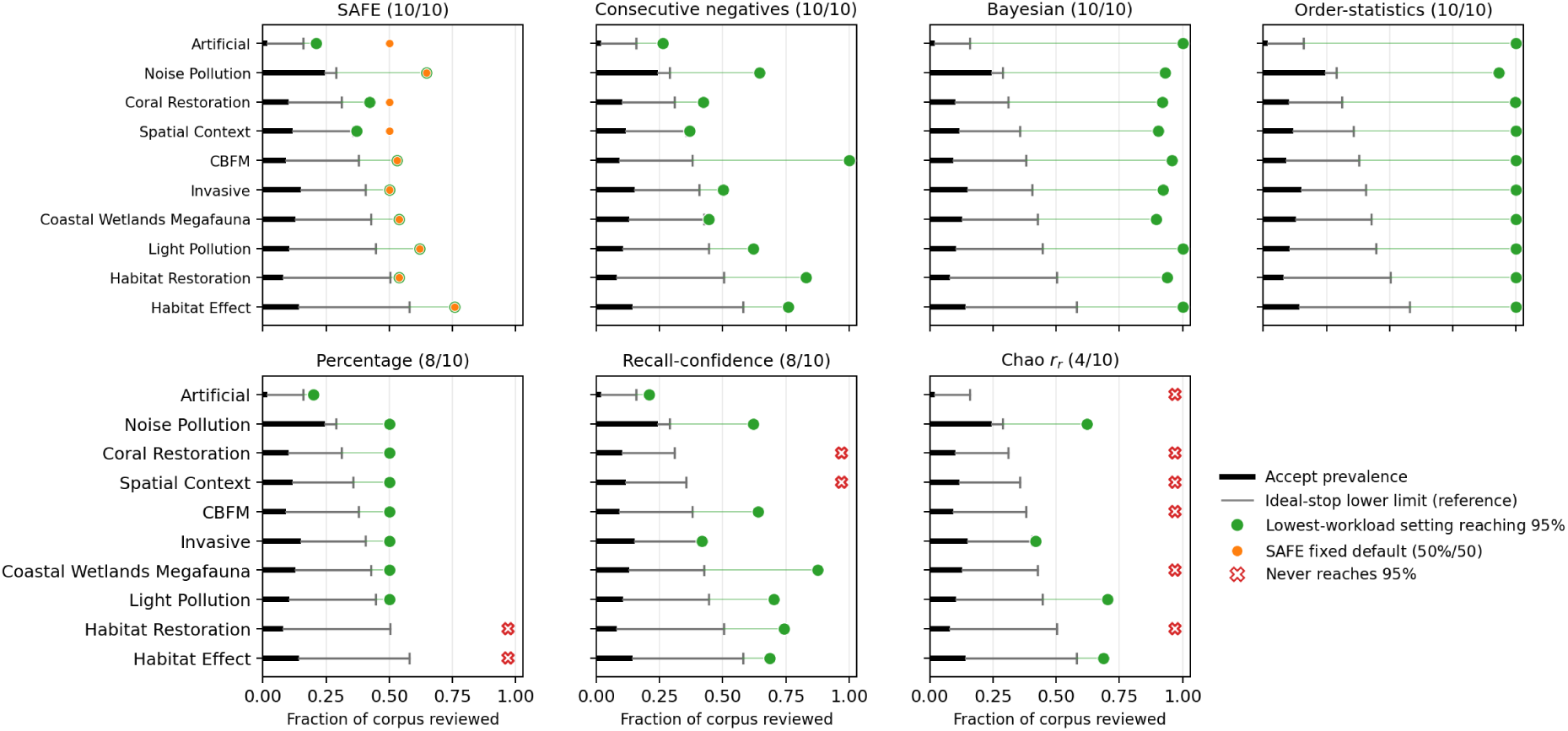
Stopping workload by review for each candidate rule, under the universal ranking, with one panel per rule. The black bar represents each review’s accept prevalence; the grey line segment runs from the end of that bar out to the ideal stopping point’s workload-to-95%-recall, which is the reference lower limit based on the universal ranking for that review. A green dot is the lowest workload at which the rule reaches 95% recall on that review; an open red cross marks reviews where the rule never reaches 95%. On the SAFE panel, an orange dot additionally marks the label-free fixed default we recommend (minimum coverage 50%, run length 50); because SAFE’s stopping point depends on which records fall in its 200-record spot-check, both SAFE markers are averaged over those draws (Supplement H).

##### Which stopping rule should a reviewer use?

We compared the seven candidate rules review by review in Figure 5, and report their cross-review aggregates in Table S7, with the full rule-by-rule breakdown in Supplement G. It is important to keep in mind that these results reflect the best-case upper bounds on what a rule family can deliver and are not what a reviewer choosing a setting in advance would obtain. Of the seven, *SAFE* was the most reliable, reaching the 95% target on all ten reviews at a mean workload of 51% (90% confidence interval over reviews [0.35, 0.66]), which was only a 12-percentage-point gap above this lower limit. The runner-up, *consecutive negatives*, also reached the 95% target on all ten and read more of the corpus (59% workload). *Bayesian* and *order-statistics* were excessively conservative, reaching the target on every review, but only by reading essentially the entire corpus (95% and 99%). The *percentage* rule had the lowest mean workload of any rule (46%) but reached 95% recall on only 8*/*10 reviews, because a fixed-fraction cutoff under-shoots when relevant records are scattered deep in the ranking. The two confidence-based rules (*recall-confidence*, *Chao r_r_*) reached target on 8*/*10 and 4*/*10, respectively, at 61% and 61% workload. (We also tested a family of “model-aware” rules that use the LLM score for every record, although none beat SAFE on reliability, and we report them in Supplement Q.)

##### A SAFE setting reviewers can pick in advance

Whilst SAFE performed best in an idealised setting (with parameters tuned to each dataset), we found that a single SAFE setting fixed in advance, with no prior knowledge of the corpus, recovered ≥ 95% of relevant records on all ten reviews while screening just 56% of the corpus on average (Table S8). This fixed setting missed 1.8% of relevant records on average and at most 3.9% on any single review, comfortably inside the standard 5% allowance. Its 56% workload is only about five percentage points more than the 51% that per-review strategies reach in hindsight.

## Discussion

We show that across ten systematic reviews spanning nearly 20,000 studies, a four-LLM ensemble selected without access to any review’s labels ranked relevant studies nearly as well as the best ranking achievable with full knowledge of each review’s annotations, so a workflow built without label-tuning can generalise across reviews without bespoke model selection for each new corpus. Applying a SAFE stopping rule chosen in advance to this ranking recovered at least 95% of relevant records on every review, missing 1.8% of relevant records on average against the 5% its target permits, while reducing the human screening workload by nearly half (56% of the corpus screened on average). A fully open-source, consumer-hardware workflow can therefore substantially reduce screening burden while maintaining the high sensitivity that systematic reviews require.

### Comparison with related work

The closest comparator to our universal-strategy result is Akinseloyin et al. (2026), who run a proprietary three-LLM ensemble that averages the models’ relevance scores (*GPT-4o Mini*, *Claude 3 Haiku*, *Gemini 1.5 Flash*) on 28 medical reviews from CLEF eHealth 2019. Their best ranking reaches mean AP 0.462 on intervention reviews and 0.341 on diagnostic-test-accuracy reviews; under an idealised stopping point, this corresponds to screening roughly 32–39% of the corpus to recover 95% of the relevant records. Our open-source four-LLM ensemble at 4-bit quantisation reaches mean AP 0.657 across ten marine-ecology reviews, with the same idealised stopping point sitting at 39% workload. Their analysis stops at the ranking layer; ours adds a stopping rule a reviewer can actually run (Step 2).

Two papers using proprietary frontier LLMs contextualise the binary accept/reject numbers in Supplement I. Fagerberg et al. (2025) report 99.7% sensitivity and 49.3% specificity from a dual *GPT-5 Thinking* + *Gemini 2.5 Pro* ensemble under an inclusive “include if either model says include” rule on 16 Cochrane reviews; Cao et al. (2026) report 96.7%/97.9% from *GPT-4.1* acting as a standalone reviewer on five medical reviews (32,357 citations). Our 95%/57% at the 50-out-of-100 midpoint cutoff falls short on accuracy, but their numbers carry API costs that scale with corpus size and rest on Cochrane-curated medical criteria, and Fagerberg et al. (2025)’s near-perfect sensitivity comes with the same sensitivity-specificity trade-off we face.

Vembye et al. (2025) reach human-level recall on psychology reviews using ten *GPT-4o-mini* replicates per record with a *k*-out-of-ten inclusion threshold; their approach draws diversity from within-model replicate variation and classifies every record, while ours averages across four different open-source LLMs and applies a stopping rule. Their proposed benchmark for automated second-screening (recall ≥ 75%, specificity ≥ 80%) is one our binary 95%/57% at the 50-out-of-100 cutoff clears on recall but not specificity, supporting our use of the ensemble as an initial screen that prioritises records for human review, not as a replacement for a second human screener. Repke et al. (2026) evaluate 15 practical stopping rules on 81 multi-domain systematic reviews and report that Callaghan-Müller-Hansen (CMH) is the only rule that never stops below target. They explicitly exclude hybrid methods from their evaluation, so SAFE’s 10*/*10 on our ecology reviews and CMH’s recommendation on theirs are complementary.

Madeyski et al. (2026) argue that summary metrics like AP and *F*_1_ can mislead when missing a relevant record costs much more than including an irrelevant one (the typical screening regime), and recommend Lost Evidence (1 − recall) and a weighted Matthews correlation coefficient as primary metrics. We use AP for the ranking comparisons but report recall and workload directly for stopping, which translate to their preferred framing without further computation. Huotala et al. (2025) benchmark nine LLMs across 24 software-engineering reviews and report that differences in screening accuracy between reviews exceed differences between LLMs, a pattern echoed by the spread in our per-review best achievable AP (0.47–0.98).

Against *ASReview* (van de Schoot et al., 2021), the established active-learning tool, our ranking reached the 95% target at lower workload on most reviews. We ran ASReview in its default configuration, tuned by its developers on a general benchmark rather than ecological corpora, so domain-specific tuning could narrow the gap, though such tuning depends on labelled data that our ensemble does not require. The two methods trade off differently: ASReview learns from the reviewer’s own decisions as screening proceeds, whereas our ensemble works from the written inclusion criteria and can rank a whole corpus before any screening begins, and can be regenerated from the same criteria and models rather than depending on which studies a reviewer marked and in what order. Combining the two is a useful future direction.

Whilst we developed the approach using ten reviews most familiar to us from the marine domain, our two out-of-domain reviews show that the method works on non-marine case studies and also matches the proprietary frontier models of their source studies (Nawrath et al., 2026; Nykvist et al., 2025). Further, our method does so without per-record API cost or sending records to a third party.

### Limitations and scope

Several limitations bear on these results and on whether to implement the method. The AP values we report are conditional on the screening prompt, the criteria wording, and the 4-bit quantised inference, though our sensitivity analyses show the ranking holds under small deviations from these settings.

We set a 95% recall target because it is standard in these workflows, but that means roughly 5% of relevant records may be missed. Our blind re-audit suggests these may be only loose-fitting records a human reviewer would keep ‘just to be sure’, but for some use-cases the risk may be intolerable, and such reviewers would be well-served by more conservative settings on the choices presented here (number of LLMs, replicates, stopping rules).

Because the ensemble reads only titles and abstracts against the screening criteria, the ranking is only as good as the criteria it is given. Where those criteria are vague, contested, or still moving, as they often are when screening begins, it will likely be misled. Small pilot screens, routine in multi-reviewer studies, may help mitigate these failures.

SAFE reached the 95% target on all ten reviews, but this is an empirical record, not a definitive baseline. On a review whose score distribution is unlike ours, using these settings could perform differently. Whilst we have provided evidence that two out-of-sample corpora perform similarly to our collection, further empirical observations will be needed to build trust in such a rule.

### Practical guidance for LLM-assisted screening

To transfer our findings to a new ecological systematic review, we provide a three-step workflow for reviewers here.

#### Step 1: Use a four-LLM mean ensemble

The specific subset {*gemma3*, *gpt-oss*, *mistral-small3*, *qwen3*} maximised mean AP across our ten reviews, and the same subset was selected on every LOO split. As shown above, ensemble size matters more than composition once three or four capable open-source LLMs are in the mix, with the added AP shrinking to within replicate-to-replicate noise by the fourth model. A reviewer with a different set of open-source LLMs on hand can substitute their own: pick any three or four comparably-sized LLMs and average all the resulting scores. For a single production screen one replicate per LLM is likely enough, since it recovers almost all of the ranking and stopping performance. However, some reviewers may find it is worth it to run replicates to obtain a variance estimate.

#### Step 2: Stop with SAFE at minimum-coverage 50%, run-length 50

This is the lowest-workload advance-selectable SAFE setting that achieved the 95% recall target on every review across 500 random spot-check draws. Mean workload across the ten reviews was 56% of the corpus (90% confidence interval [0.50, 0.66]); it missed 1.8% of relevant records on average (mean recall 0.982) and at most 3.9% on any single review (lowest recall 0.961), inside the 5% the 95% target permits. A reviewer who can afford a small labelled subset for tuning can recover roughly five percentage points by tuning per-review, but choosing tighter SAFE values without labels exposes the workflow to failure modes documented in Supplement H.

#### Step 3: Treat the LLM as an initial screen instead of a replacement screener

The full workflow uses the ensemble to *rank* records and SAFE to decide *when to stop reading*; humans still screen everything above the stopping point. Asked to make accept/reject decisions on its own (using a 50-out-of-100 cutoff on the averaged relevance score), the LLM ensemble agrees with the human review leads less often than two human screeners typically agree with each other. Chance-corrected agreement (Cohen’s *κ*) is about 0.24, well below the ≈ 0.8 mean *κ* reported for human reviewers screening abstracts (Hanegraaf et al., 2024) (Supplement I). Used as a triage layer this matters less than it may appear, because SAFE sets a 95% recall minimum on relevant records, so the dominant remaining source of decision error is the human screener’s own variability, not the LLM’s. Used as a binary classifier, though, the same numbers warn against replacement, as a reviewer who swaps human screening for the LLM at any single threshold should expect substantially more disagreement than a second human would give, so we do not recommend it.

The full pipeline (four-LLM ensemble, SAFE) uses LLMs small enough to run on a high-end consumer laptop, though we ran it on an NVIDIA H100 GPU for this study. For our ten reviews, wall-clock for the ensemble screen on the H100 ranged from ∼ 2 hours on the smallest corpus (*N* = 285) to ∼ 25 hours on the largest (*N* = 3,877). Once a reviewer has committed to LLM-assisted screening, the stopping-rule overhead is negligible compared with the ensemble screen itself.

To make this workflow straightforward to reproduce and to apply to new reviews, we release it as an open-source R package, screenllm (https://github.com/s-spillias/screenllm), with the recommended four-LLM mean ensemble and advance-choosable SAFE default built in, plus a companion app for screening the records above the stop point.

### Future directions

The clearest next step is to test the workflow on additional datasets, both within and beyond environmental science. Running the same rank-then-stop protocol across fields with contrasting criteria would show how generalisable these results are. A second direction concerns the ensemble itself. Our five LLMs were of comparable size and, apart from *deepseek-r1*, of similar non-reasoning design, and returns saturated by three or four models; ensembles that deliberately mix architectures or training regimes, for instance pairing reasoning and non-reasoning models, might supply diversity that similar models cannot and lift the performance ceiling we observed. The best-performing subset may also be discipline-specific, so we expect the selection procedure to transfer more reliably than any single four-model set, and confirming this across disciplines is a natural extension of this work.

## Acknowledgements

SS was funded by a CSIRO R+ Postdoctoral Fellowship. SSw was supported by the Jock Clough Marine Foundation Oceans Chair at UWA. MS was supported by an Australian Research Council (ARC) Discovery Early Career Researcher Award no. DE220100079. BW was supported by funding awarded to VK by the Department of Natural Resources and Environment Tasmania. VK was supported by an ARC Discovery Project (DP240102334, awarded to S. Swearer).

## Data Availability

The complete analysis code, including all scripts, configuration files, and analysis tools, is available at https://github.com/s-spillias/screenllm-paper. The rec ommended workflow is additionally distributed as an open-source R package, `screenllm` (https://github.com/s-spillias/screenllm), for anyone who wishes to apply the method to their own reviews.

## Author Contributions

SS: Conceptualization, Methodology, Software, Validation, Formal analysis, Investigation, Resources, Data Curation, Writing - Original Draft, Writing - Review & Editing, Visualization, Project administration. CB: Formal analysis, Writing - Original Draft, Writing - Review & Editing. VK: Conceptualization, Investigation, Data Curation, Writing - Original Draft, Writing - Review & Editing. SSw: Conceptualization, Writing - Original Draft, Writing - Review & Editing. LAT, AE, JR, MS, AT, BW: Data Curation, Writing - Review & Editing.

## Statement on the Use of Generative AI

Generative AI tools, specifically Microsoft Copilot and Claude Code, were utilized in the preparation of this manuscript to assist with language refinement and analytical code development. All scientific content, data interpretation, and conclusions were verified by the authors to ensure accuracy and integrity.

