## Supplemental Information for "Operationalising LLM-assisted screening of literature to support systematic reviews"

August 18, 2026

[1]CSIRO Environment, Hobart, Australia
[2]Centre for Marine Socioecology, University of Tasmania, Hobart, Australia
[3]Oceans Institute, University of Western Australia, Perth, Australia
[4]Global Wetlands Project, Australian Rivers Institute, School of Environment and Science, Griffith University, Gold Coast, QLD 4222, Australia
[5]Institute for Marine and Antarctic Studies (IMAS), University of Tasmania, Hobart, Australia

### Supplement A: Confidently-misplaced positives under the universal ranking

| **Review** | $N_+$ | **Score** $< 30$ | **Score** $< 20$ | **Score** $< 10$ | **Bot.** $25\%$ | **Bot.** $10\%$ |
|---|---|---|---|---|---|---|
| Artificial | 73 | 0% | 0% | 0% | 0% | 0% |
| CBFM | 101 | 0% | 0% | 0% | < 1% | 0% |
| Coastal Wetlands Megafauna | 340 | 2% | 1% | < 1% | 1% | < 1% |
| Coral Restoration | 340 | 3% | < 1% | 0% | 0% | 0% |
| Habitat Effect | 52 | 0% | 0% | 0% | 0% | 0% |
| Habitat Restoration | 228 | 1% | < 1% | < 1% | < 1% | 0% |
| Invasive | 90 | 2% | 1% | 0% | 0% | 0% |
| Light Pollution | 30 | 0% | 0% | 0% | 0% | 0% |
| Noise Pollution | 79 | 0% | 0% | 0% | 0% | 0% |
| Spatial Context | 456 | < 1% | 0% | 0% | < 1% | 0% |
| **Mean across reviews** | — | **0.9%** | **0.4%** | **0.1%** | **0.3%** | **0.0%** |

Table S1: Confidently-misplaced positives under the universal ranking. $N_+$ is the number of human-accepted (positive) records per review. The next three columns give the fraction of those positives that received an aggregated LLM score below 30, 20, and 10 on the 0–100 scale, the regime where the LLM is most confident a record should be rejected. The last two columns give the fraction that fell into the bottom quartile and bottom decile of the universal ranking. Lower is better in every column. Values are rounded to whole percent, and "$< 1\%$" marks single-record errors that round to zero. Confidently-misplaced positives were rare on every review, with a cross-review mean of under $1\%$ of human-accepted records scoring below $30$. *Habitat Effect* uses the corrected labels described in the Methods.

#### Supplement B: Does prevalence or review size predict difficulty?

We tested the obvious candidate explanations for the per-review spread in best achievable AP. We found that accept prevalence is significantly and positively correlated with best achievable AP ($r = 0.71$, $p = 0.021$, $95\%$ CI $[0.15, 0.93]$), so lower-prevalence reviews tended to rank slightly less well, whereas review size showed no detectable association ($r = -0.27$ on $\log_{10} N$, $p = 0.45$, $95\%$ CI $[-0.77, 0.43]$; Figure S1). The association is statistically significant across these ten reviews, but additional datapoints would be needed to treat this finding as conclusive. The two 'hardest' reviews, *Artificial* and *Habitat Effect*, have very different prevalences ($1.9\%$ and $14.3\%$), so difficulty also reflects the criteria themselves, in particular their compositional structure and the domain knowledge they assume.

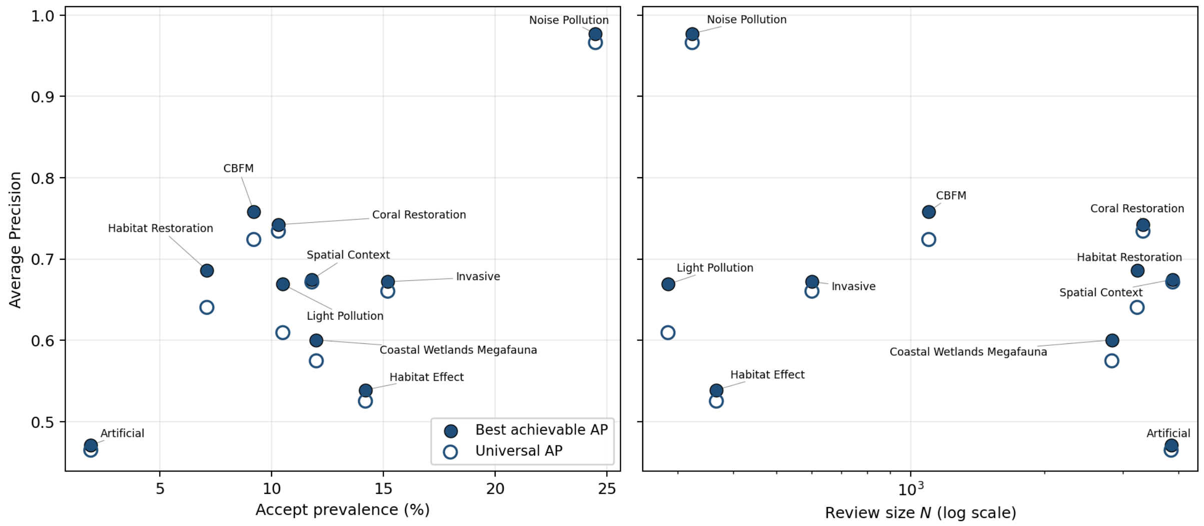


Figure S1: Per-review best achievable AP (filled circles) and AP under the universal strategy (open circles), against accept prevalence (**left**) and review size $N$ on a log scale (**right**). Accept prevalence is positively associated with best achievable AP across these ten reviews (Pearson $r = 0.71$), whereas review size is not ($r = -0.27$). However, the two lowest-AP reviews, *Artificial* and *Habitat Effect*, have markedly different prevalences, indicating that prevalence is only one of several factors contributing to review difficulty.

### Supplement C: Per-review best achievable AP and best ranking strategy

| Review | Best AP | $\Delta$AP (univ.) | Workload$_{95}$ | Aggregator | $r$ | Model subset |
|---|---|---|---|---|---|---|
| Noise Pollution | 0.977 | −0.011 | 0.29 | mean | 3 | deepseek-r1; gemma3 |
| CBFM | 0.758 | −0.034 | 0.38 | mean | 3 | deepseek-r1; gemma3; qwen3 |
| Coral Restoration | 0.742 | −0.008 | 0.31 | mean | 3 | deepseek-r1; gemma3; mistral-small3; qwen3 |
| Habitat Restoration | 0.686 | −0.046 | 0.50 | mean | 3 | gpt-oss; qwen3 |
| Spatial Context | 0.675 | −0.003 | 0.36 | mean | 2 | gemma3; gpt-oss; mistral-small3; qwen3 |
| Invasive | 0.672 | −0.012 | 0.41 | mean | 3 | gpt-oss; mistral-small3 |
| Light Pollution | 0.669 | −0.060 | 0.45 | mean | 3 | gemma3; mistral-small3 |
| Coastal Wetlands Megafauna | 0.600 | −0.026 | 0.43 | mean | 3 | deepseek-r1; gpt-oss; mistral-small3 |
| Habitat Effect | 0.539 | −0.014 | 0.58 | median | 3 | gpt-oss; qwen3 |
| Artificial | 0.471 | −0.006 | 0.16 | mean | 3 | gpt-oss; mistral-small3; qwen3 |

Table S2: Per-review with-labels best achievable AP: highest mean AP achievable across replicate combinations, and the strategy that achieves it. $r$ is the number of replicates per LLM used by the ensemble. **$\Delta$AP (univ.)** is the per-review gap from this best achievable AP to the universal four-LLM ensemble. **Workload$_{95}$** is the ideal-stop workload-to-95%-recall under the universal ranking (the *precalc* ideal stopping point, i.e. the lowest workload any rule could reach at 95% recall on that ranking, computed using the true accept count); we report it on the same row as Best AP to flag that a high best achievable AP does not imply low screening workload, since AP and workload-to-recall summarise different parts of the ranking.

### Supplement D: Error propagation

The ranking and stopping outcomes were calculated based on two inputs with known variance: stochastic LLM scores and human screening labels. To quantify uncertainty, we propagated both sources through universal-ensemble Average Precision (AP), ideal-stop workload, recall at a fixed $55\%$ of the corpus (a conservative proxy for the advance-fixed SAFE default of $56\%$), and the corresponding miss rate ($1-\mathrm{recall}$). We used Monte Carlo resampling with $B = 1000$ draws per review. Each draw preserved the score distribution underlying the reported per-review AP, ensuring comparability between resampled metrics and point estimates. Metrics were recomputed for each review and averaged across reviews. Label uncertainty was estimated from a stratified re-audit of three reviews, each independently re-screened by two assessors, and the resulting stratum-specific error rates were applied corpus-wide.

**Replicate variance.** To estimate uncertainty arising from model stochasticity, each draw resampled the three replicates of each of the four universal models and recomputed all metrics. Two resampling models were evaluated. The nonparametric bootstrap resampled the three replicates with replacement and averaged the resulting scores. The parametric approach sampled scores from a $t$ distribution with two degrees of freedom, centred on the replicate mean with scale equal to the replicate standard error and truncated to $[0, 100]$. The low degrees of freedom reflect uncertainty in estimating variance from only three replicates. Both approaches produced similar results. The cross-review mean AP interval was $[0.651, 0.658]$ under the bootstrap and $[0.646, 0.656]$ under the parametric model. Replicate-induced uncertainty was small across all reviews (approximately 0.01 AP). The widest per-review interval occurred for *Habitat Restoration*, spanning $[0.634, 0.645]$ under the bootstrap and $[0.622, 0.644]$ under the parametric model. These results indicate that ensemble rankings are stable across stochastic reruns.

**Label uncertainty from a stratified re-audit.** Human screening labels represent a second source of uncertainty. Previous re-review work assessed only records where the ensemble and original screener disagreed, leaving error rates among concordant records unmeasured. To estimate corpus-wide label uncertainty, we conducted a stratified, blinded re-audit across all ten reviews. Records were assigned to one of five strata defined by the original label and ensemble-score band: *missed-positive* (ensemble $< 30$, human Accept), *contested* (ensemble $\geq 70$, human Reject), two high- and low-confidence agreement strata, and a *mid* stratum (ensemble score 30–70). Re-screeners were provided only with title, year, journal, abstract, and review inclusion criteria. They were blinded to ensemble scores, original decisions, and stratum assignments. Records were randomly interleaved across strata. The re-audit covered *Light Pollution*, *Spatial Context*, and *Coral Restoration*, each independently assessed by two readers. Error-rate estimates were pooled across reviews. The *missed-positive* stratum was evaluated separately through targeted adjudication because records defined by disagreement cannot be assessed blindly. For each record, reviewers determined whether it represented a genuine ensemble miss or an original over-inclusion. Because strata were sampled at unequal rates, corpus-level error rates were estimated as

$$E = \sum_h \frac{N_h}{N} e_h,$$

where $e_h$ is the stratum-specific error rate and $N_h/N$ its corpus prevalence.

**Propagation of label uncertainty.** We propagated label uncertainty by replacing a single global correction rate with stratum-specific error rates estimated from the audit. For each Monte Carlo draw, stratum correction probabilities were sampled from Beta posteriors defined by the audit counts, labels were perturbed accordingly, and all metrics were recomputed. The audit revealed substantial asymmetry across strata. Disagreement with the original labels was common within the *confident-accept* stratum but rare within the *confident-reject* stratum. Because title-and-abstract screening uses an intentionally inclusive acceptance criterion, we interpreted the former as variation in screening threshold rather than mislabelling. Consequently, confident-accept disagreements were not treated as label errors in the primary analysis. Uncertainty propagation therefore included the *contested* stratum and the small measured error rates in the remaining strata. As a conservative sensitivity analysis, we also considered an adversarial scenario in which all disagreements, including those in the confident-accept stratum, were treated as corrected labels.

**Propagated Error.** Across the three re-audited reviews, the estimated error rate in the *contested* stratum was 20%, although substantial reviewer-to-reviewer variability was observed. One reviewer supported the ensemble on 36% of contested records, whereas the other did so on 3%, with review-specific rates ranging from 5% to 30%. Estimated error rates in the confident-reject and mid strata were 0.5% and 0.7%, respectively. The confident-accept stratum showed a 46% disagreement rate, consistent with differences in screening threshold rather than systematic labelling error. After reweighting by stratum prevalence, the corpus-level disagreement rate was approximately 7%. All 11 records in the missed-positive stratum were judged irrelevant by both re-reviewers, indicating original over-inclusion rather than genuine ensemble misses. Under the primary uncertainty model, the cross-review mean AP increased from the point estimate of 0.657 to a label-adjusted interval of $[0.66, 0.69]$. The combined replicate-and-label interval was also $[0.66, 0.69]$, reflecting the small contribution of replicate variance. Recall at the advance-fixed SAFE workload remained at least 0.82 for every review, and the cross-review mean miss rate remained within $[0.02, 0.05]$. Under the adversarial sensitivity analysis, mean AP decreased to $[0.36, 0.42]$ and the minimum recall to 0.79. We interpret this scenario as a conservative lower bound rather than a realistic estimate.

| Review | AP | Replicate 90% CI | Label-adjusted range |
|---|---|---|---|
| Noise Pollution | 0.966 | $[0.964, 0.969]$ | $[0.892, 0.939]$ |
| Coral Restoration | 0.734 | $[0.731, 0.735]$ | $[0.712, 0.801]$ |
| Habitat Restoration | 0.640 | $[0.634, 0.645]$ | $[0.618, 0.698]$ |
| CBFM | 0.724 | $[0.713, 0.726]$ | $[0.677, 0.788]$ |
| Spatial Context | 0.672 | $[0.668, 0.676]$ | $[0.642, 0.690]$ |
| Invasive | 0.660 | $[0.648, 0.664]$ | $[0.668, 0.755]$ |
| Light Pollution | 0.609 | $[0.576, 0.623]$ | $[0.580, 0.704]$ |
| Coastal Wetlands Megafauna | 0.575 | $[0.569, 0.577]$ | $[0.604, 0.675]$ |
| Habitat Effect | 0.525 | $[0.515, 0.535]$ | $[0.535, 0.650]$ |
| Artificial | 0.465 | $[0.430, 0.473]$ | $[0.366, 0.503]$ |
| **Mean** | **0.657** | $[0.651, 0.658]$ | $[0.662, 0.693]$ |

Table S3: Propagation of replicate and label uncertainty in universal-ensemble Average Precision (AP). Point estimates are based on the original labels. Replicate intervals reflect bootstrap resampling of model replicates, and label-adjusted intervals reflect Monte Carlo propagation of stratum-specific label-error rates estimated from the blinded re-audit. Error rates were pooled across audited reviews and applied corpus-wide.

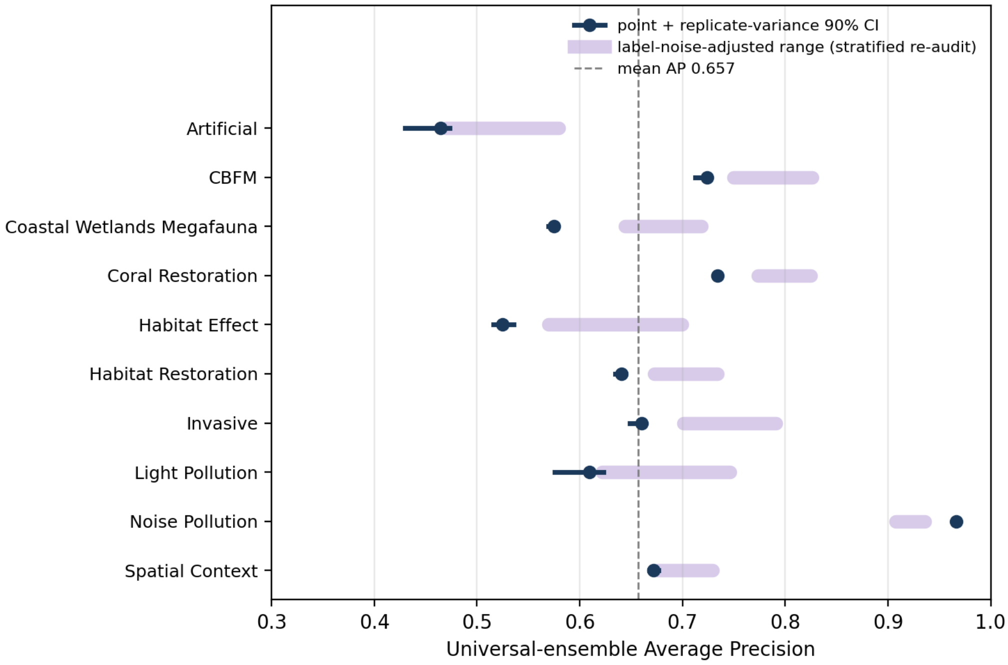


Figure S2: Per-review universal-ensemble Average Precision (AP) with propagated uncertainty. Points denote AP estimates; dark bars show 90% replicate-variance intervals and light bars show label-adjusted intervals derived from re-audit error rates. Label uncertainty contributes more to interval width than replicate variance across all reviews.

### Supplement E: Sensitivity to workflow choices

Beyond the choice of which LLMs to run (the ensemble-size and composition analysis of Supplement F and the main text), a reviewer deploying this workflow makes three further design choices. These are how the screening prompt is written, how the inclusion criteria are phrased, and the numerical precision at which the LLMs are served. We tested the sensitivity of the recommended four-LLM *mean* ensemble to each of these on three reviews that span the range of ranking difficulty. *Light Pollution* has the largest universal-versus-best gap, *Noise Pollution* is the easiest to rank, and *CBFM* is a mid-difficulty review an order of magnitude larger. For every comparison we held the records, models, and replicate count fixed and changed only the one factor, then bootstrapped the three replicates on both arms ($B = 2000$), so that we report each $\Delta$AP against the same replicate-to-replicate noise ($\approx 0.01$ AP) that bounds the main results. A $90\%$ interval that excludes zero marks a factor the ranking is genuinely sensitive to. Because we compute the absolute AP on each arm from these matched re-runs, it can differ by a few thousandths from the point estimates in Table S3, but the within-comparison $\Delta$AP is unaffected.

**Prompt design and criteria phrasing.** Table S4 evaluates the robustness of the recommended four-LLM mean ensemble to prompt formulation and criteria wording. Binary include/exclude prompts reduced ranking performance across reviews, whereas rewording the eligibility criteria had negligible effect. A single holistic relevance score improved AP only for *Light Pollution*, the most challenging review, suggesting that holistic scoring may better capture relevance when eligibility depends on overall judgement rather than criterion-specific evidence.

| **Review** | **Factor / variant** | **Base AP** | **Var. AP** | **ΔAP** | **90% CI** |
|---|---|---|---|---|---|
| Light | Prompt: holistic single score | 0.599 | 0.723 | $+\mathbf{0.123}$ | $[0.103, 0.160]$ |
| Light | Prompt: binary 0/100 | 0.599 | 0.556 | $-\mathbf{0.043}$ | $[-0.081, -0.018]$ |
| Light | Criteria: reworded | 0.599 | 0.581 | $-0.018$ | $[-0.032, 0.017]$ |
| Noise | Prompt: holistic single score | 0.968 | 0.969 | $+0.001$ | $[-0.004, 0.004]$ |
| Noise | Prompt: binary 0/100 | 0.968 | 0.947 | $-\mathbf{0.021}$ | $[-0.053, -0.019]$ |
| Noise | Criteria: reworded | 0.968 | 0.966 | $-0.002$ | $[-0.006, 0.000]$ |
| CBFM | Prompt: holistic single score | 0.753 | 0.766 | $+0.013$ | $[-0.004, 0.022]$ |
| CBFM | Prompt: binary 0/100 | 0.753 | 0.719 | $-\mathbf{0.034}$ | $[-0.054, -0.033]$ |
| CBFM | Criteria: reworded | 0.753 | 0.745 | $-0.008$ | $[-0.031, -0.007]$ |

Table S4: Sensitivity of the recommended four-LLM mean ensemble to prompt formulation and criteria wording. Average Precision (AP) under each variant is compared with the standard per-criterion partial-credit prompt. Differences in AP (ΔAP) and their 90% bootstrap confidence intervals were estimated from replicate resampling ($B = 2000$). Variants comprised a holistic relevance score, a binary include/exclude decision, and a reworded version of the eligibility criteria. Binary prompting consistently reduced performance, whereas criteria rewording had negligible effect.

**Quantisation.** The three quantisable models were evaluated at 8-bit (*Q8_0*) and 16-bit (*fp16*) precision on the same three reviews, with *gpt-oss* held at MXFP4 because alternative precision formats were unavailable. Ensemble performance was highly stable across precisions (Table S5), with no change in AP exceeding 0.011 on any review.

| **Review** | $N$ | **Prev.** | **AP (Q4)** | **AP (Q8): $\Delta$ [$90\%$ CI]** | **AP (fp16): $\Delta$ [$90\%$ CI]** |
|---|---|---|---|---|---|
| Light | 285 | 10.5% | 0.599 | 0.597: $-0.002$ $[-0.028, 0.033]$ | 0.590: $-0.009$ $[-0.033, 0.028]$ |
| Noise | 323 | 24.5% | 0.968 | 0.957: $\mathbf{-0.011}$ $[-0.014, -0.009]$ | 0.962: $\mathbf{-0.006}$ $[-0.012, -0.004]$ |
| CBFM | 1,097 | 9.2% | 0.753 | 0.749: $-0.004$ $[-0.017, 0.010]$ | 0.744: $-0.009$ $[-0.019, 0.002]$ |

Table S5: Sensitivity of ensemble Average Precision (AP) to model quantisation. The three quantisable models were evaluated at 8-bit (Q8_0) and 16-bit (fp16) precision, with *gpt-oss* fixed at MXFP4. Differences in AP relative to the production configuration were small across all reviews, indicating that ensemble ranking performance is largely insensitive to numerical precision.

### Supplement F: Ensemble size and replicate count, review by review

**Replicate count.** Increasing from one to three replicates per model improved cross-review mean AP by only 0.008 (Table S6), comparable to the replicate-level variability ($\approx 0.01$ AP) and with negligible change in stopping workload or recall, and the same flatness holds review by review (Figure S3); additional replicates mainly provide variance estimates (Supplement D).

Table S6: Sensitivity of ensemble performance to the number of replicates per model. Results are shown for the recommended four-LLM mean ensemble as cross-review averages on the deduplicated corpus. For $r < 3$, metrics were averaged across all replicate combinations. Average Precision (AP), workload required to achieve 95% recall ($\text{Workload}_{95}$), and recall at a fixed $55\%$ workload ($\text{Recall}_{55}$, a conservative proxy for the advance-fixed SAFE default) show minimal improvement beyond a single replicate, indicating rapidly diminishing returns from additional sampling.

| $r$ | AP | $\text{Workload}_{95}$ | $\text{Recall}_{55}$ |
|---|---|---|---|
| 1 | 0.649 | 0.390 | 0.978 |
| 2 | 0.655 | 0.387 | 0.978 |
| 3 | 0.657 | 0.385 | 0.978 |

**Ensemble size.** On every review the best achievable AP rises from one LLM to two or three and then flattens or dips slightly, so a larger ensemble is never reliably better (Figure S4).

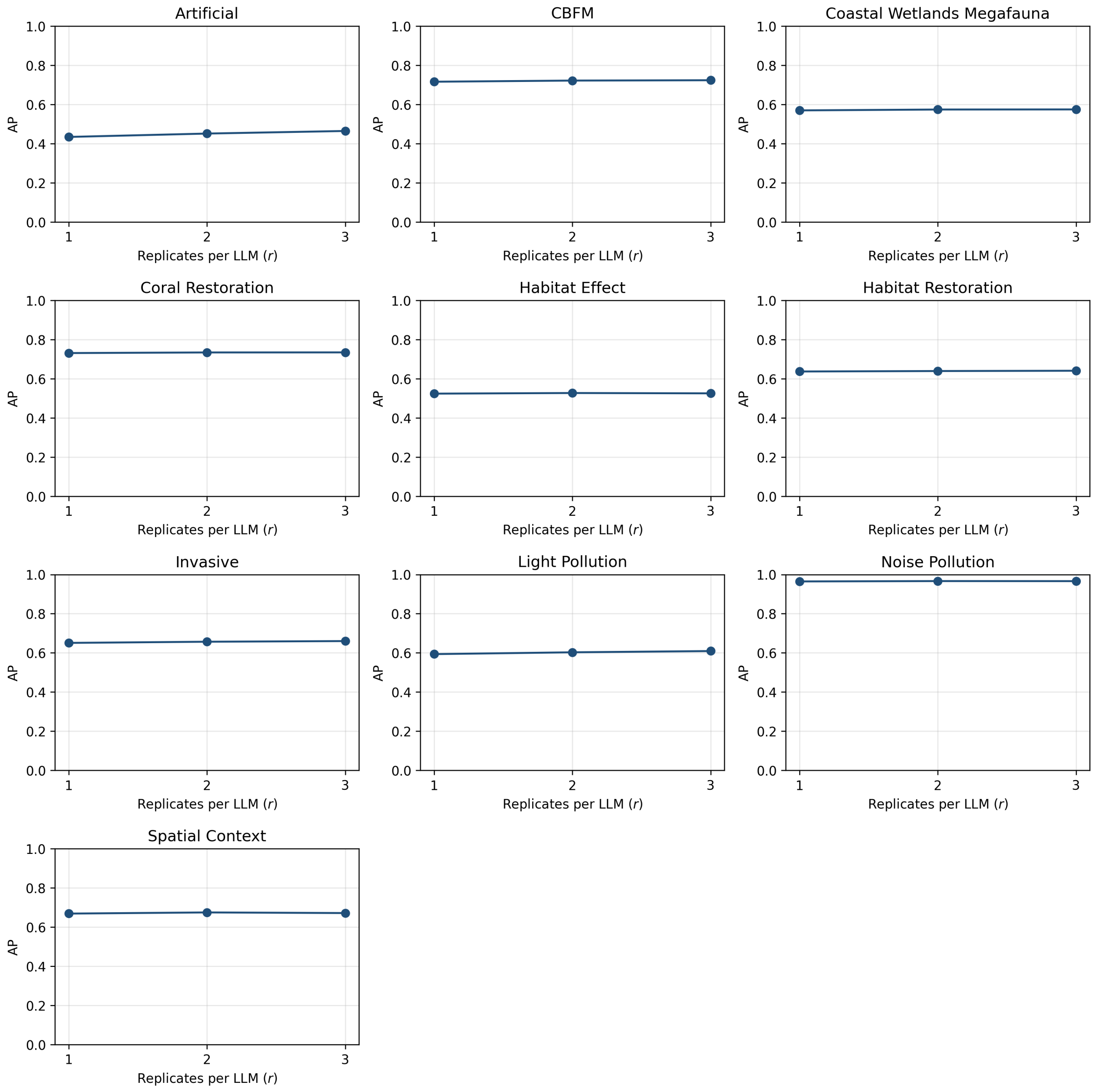


Figure S3: For each review, Average Precision under the recommended four-LLM *mean* ensemble at each replicate count $r \in \{1, 2, 3\}$, averaged across replicate combinations as in Table S6.

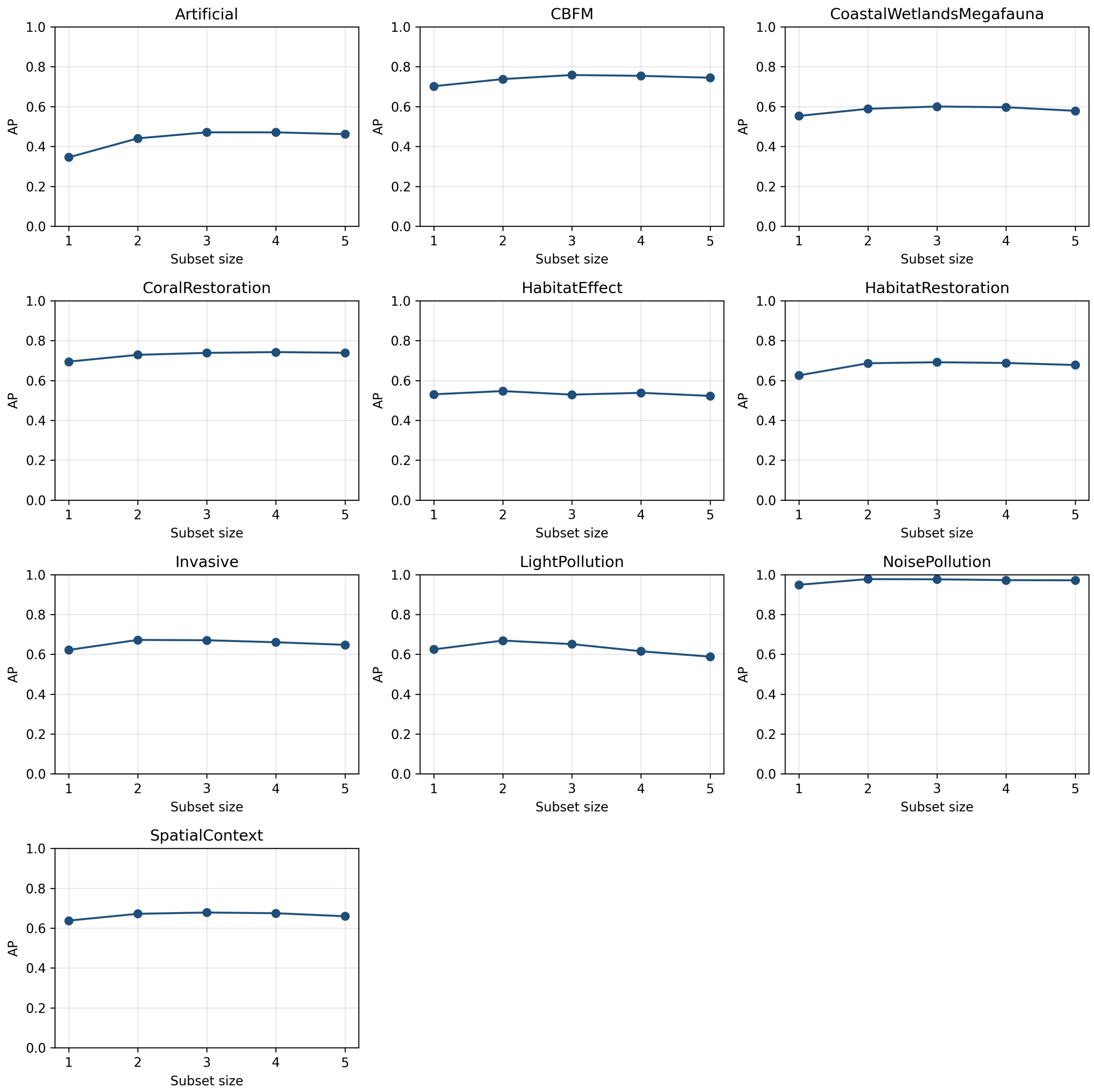


Figure S4: For each review, the best AP achievable at each ensemble size $s \in \{1, \ldots, 5\}$. Lines are typically not strictly increasing. On most reviews, allowing a fifth LLM leaves the best AP unchanged or slightly lower, and the per-review differences are within the replicate-to-replicate AP variability of the ensemble.

### Supplement G: Stopping-rule comparison

| Stopping rule | Reviews hitting target | Mean workload | Median workload | Mean recall |
|---|---|---|---|---|
| *Candidate rules* | | | | |
| **SAFE** | **10/10** | **0.51** | **0.52** | **0.974** |
| Consecutive negatives | 10/10 | 0.59 | 0.56 | 0.981 |
| Bayesian | 10/10 | 0.95 | 0.93 | 1.000 |
| Order-statistics | 10/10 | 0.99 | 1.00 | 1.000 |
| Percentage | 8/10 | 0.46 | 0.50 | 0.980 |
| Recall-confidence | 8/10 | 0.61 | 0.66 | 0.989 |
| Chao $r_r$ | 4/10 | 0.61 | 0.65 | 0.986 |
| Pre-calculated (ideal stopping point) | 10/10 | 0.39 | 0.39 | 0.956 |

Table S7: Stopping-rule performance on universal rankings. Candidate stopping rules are shown in the upper block and the ideal-stopping-point reference in the bottom row. The ideal stopping point assumes knowledge of the true number of relevant records and is therefore reported only as a lower bound on achievable workload. For *Consecutive negatives*, *Percentage*, and *SAFE*, results correspond to the lowest-workload parameter setting that achieved at least 95% recall on each review within the search grids defined in Methods. "Reviews hitting target" reports the number of reviews for which at least one parameter setting achieved the recall target. Workload and recall are summarised across the selected per-review settings. Because parameter values were chosen retrospectively using review labels, these results represent optimistic upper bounds on achievable performance. The advance-fixed SAFE configuration (Table S8) provides the deployable evaluation using a pre-specified setting. Model-aware stopping rules are reported separately in Supplement Q.

#### Supplement H: SAFE robustness to spot-check noise

| Min. coverage | Run length | Reviews hit at $\geq 95\%$ draws | Mean workload | Mean recall | Notes |
|---|---|---|---|---|---|
| 50% | 50 | **10/10** | **0.56** | **0.982** | Recommended advance-fixed default |
| 50% | 100 | 10/10 | 0.68 | 0.989 | Higher workload, no recall gain |
| 5–50% | 250 | 10/10 | 0.81–0.83 | $\sim 0.996$ | Same family at longer run length, conservative on workload |
| 20–25% | 100 | 8/10 | 0.60 | 0.979 | Misses target on 2 reviews |
| 20–25% | 50 | 7/10 | 0.44–0.45 | 0.96 | Lower coverage, misses target on 3 reviews |
| 5–10% | 100 | 7/10 | 0.59 | 0.975 | Misses target on 3 reviews |
| 5–10% | 50 | 6/10 | 0.42 | 0.943 | Smallest minimum coverage and lowest workload, but misses target on 4 reviews |

Table S8: Advance-choosable SAFE settings (minimum-coverage fraction and consecutive-negative run length), evaluated as a fixed setting (no per-review tuning) across 500 independent 200-record spot-check draws per review. "Reviews hit at $\geq 95\%$ draws" counts reviews where SAFE met the 95% recall target on at least 95% of the 500 draws. The run-length-250 row collapses five minimum-coverage values that produced essentially identical workload, because the consecutive-negatives gate dominates at long run lengths. The recommended default (50% minimum coverage, run length 50) is the unique advance-fixed setting that reaches 10/10 at the lowest workload.

**Why a noisy spot-check does not move SAFE's stopping point.** The spot-check estimate of the positive count varied substantially across our 500 draws per review. On *Artificial*, for example, it ranged from 19 to 96 against a true count of 73. This variation enters one of SAFE's internal criteria, yet the resulting workload changed little. The stability follows from SAFE's structure, which requires three conditions to be satisfied simultaneously before it stops. The minimum-coverage criterion is satisfied once a chosen fraction of the corpus has been screened. The consecutive-negatives criterion is satisfied once the last $L$ records are all negative. The spot-check criterion is satisfied once the relevant records found number at least twice the spot-check's estimate of the corpus total. SAFE stops only when the last of these three conditions is met. At each review's tuned settings that final condition is always the consecutive-negatives criterion, so the spot-check criterion has already been satisfied by the time SAFE stops, and a different spot-check draw does not change the stopping point. Pre-calculated rules that set their target directly from a single spot-check have no such redundancy, so any noise in the spot-check shifts their stopping point with it.

**Settings where SAFE can fail.** To confirm that SAFE does respond to spot-check noise under other parameters, we re-ran the same bootstrap at settings the per-review tuning had rejected. At these settings the spot-check criterion becomes the limiting condition, and the workload varies considerably. At a $5\%$ minimum-coverage, $50$-run-length setting on *Habitat Restoration*, the $500$ draws gave a workload SD of about $0.11$ and missed the $95\%$ target on every draw, and on *Coral Restoration* the same setting reached the target on only $1\%$ of draws. Spot-check noise is therefore a genuine risk for SAFE, but it has no influence at the settings each review's tuning selects. A reviewer using SAFE without label-aware tuning is exposed to these failure modes, which makes the choice of SAFE settings the component of the workflow most sensitive to mis-specification.

### Supplement I: LLM–human classification agreement

The workflow we recommend uses the LLM ensemble as a ranking-and-triage tool. A human screens every record above the SAFE stopping point, and the pipeline excludes the rest, so the LLM never makes the final accept or reject decision. To put that choice in context, this supplement reports how well the ensemble would agree with the human review leads if it did make those decisions on its own. That is the number a practitioner needs to judge how far the ensemble could replace, rather than complement, a human screener.

For each review we treated the universal-strategy ensemble as a standalone screener. It "accepts" a record when its averaged $0$–$100$ score is at least $50$, the natural midpoint of the scale, and "rejects" otherwise. We then compared that decision against the human review leads' decision, record by record. Cohen's $\kappa$ is the standard chance-adjusted measure of how much two raters agree beyond chance, where $0$ is chance-level agreement and $1$ is perfect agreement. Widely cited bands are $0$–$0.2$ slight, $0.21$–$0.40$ fair, $0.41$–$0.60$ moderate, and $0.61$–$0.80$ substantial (Landis and Koch, 1977). Across the ten reviews the mean $\kappa$ between the ensemble's decision and the human label was $0.24$ ($90\%$ CI $[0.16, 0.32]$, range $0.07$–$0.56$). The ensemble agreed with the humans on $62\%$ of records, captured $95\%$ of the records humans accepted (sensitivity), and correctly excluded $57\%$ of those they rejected (specificity, Table S9). At this midpoint cutoff it reached $\kappa \geq 0.40$ on two reviews (*Coral Restoration*, $\kappa = 0.56$, and *Coastal Wetlands Megafauna*, $\kappa = 0.42$) and fell below $0.20$ on six.

The ensemble is not a poor screener overall. Its mean AP across the ten reviews is $0.67$, so the ranking it produces is informative. What is conservative is the binary accept-or-reject decision at a fixed $50$-point cutoff, consistent with our broader finding that a score threshold does not transfer across reviews (Supplement K). The practical message is simple. A reviewer hoping to replace a human screener with the ensemble's decision at any single threshold should expect agreement well below what a second human reviewer would provide. Hanegraaf et al. (2024) go further and propose $\kappa \geq 0.8$ as a reasonable acceptance target for machine-assisted screening, matching typical human inter-reviewer agreement; the ensemble's standalone $\kappa$ of $0.24$ falls well short of that bar. A reviewer using the ensemble as a triage layer, screening only the records above SAFE's stopping point, operates in a different regime, where SAFE sets a $95\%$ recall minimum and the dominant remaining source of error is the human screener's own variability, not the LLM's.

| **Review** | $N$ | **Prev.** | **Sens.** | **Spec.** | **Agree.** | $\kappa$ |
|---|---|---|---|---|---|---|
| Coral Restoration | 3315 | 10.3% | 0.90 | 0.89 | 0.89 | 0.56 |
| Coastal Wetlands Megafauna | 2628 | 12.9% | 0.83 | 0.80 | 0.80 | 0.42 |
| Invasive | 597 | 15.1% | 0.96 | 0.69 | 0.73 | 0.38 |
| Habitat Effect | 364 | 14.3% | 0.88 | 0.63 | 0.67 | 0.28 |
| Spatial Context | 3877 | 11.8% | 0.99 | 0.49 | 0.55 | 0.18 |
| Habitat Restoration | 2827 | 8.1% | 0.96 | 0.56 | 0.59 | 0.16 |
| Noise Pollution | 322 | 24.5% | 1.00 | 0.27 | 0.45 | 0.15 |
| Light Pollution | 285 | 10.5% | 0.97 | 0.37 | 0.43 | 0.10 |
| Artificial | 3844 | 1.9% | 0.99 | 0.74 | 0.74 | 0.09 |
| CBFM | 1096 | 9.2% | 0.99 | 0.31 | 0.37 | 0.07 |
| **Mean** | | | 0.95 | 0.57 | 0.62 | **0.24** |

Table S9: Standalone classification agreement between the universal-strategy LLM ensemble (decision: $A \geq 50$ on the $0$–$100$ aggregated score) and the human review leads, per review. Sensitivity and specificity are with respect to the human label. Cohen's $\kappa$ sits well below the $\approx 0.8$ mean reported for human inter-reviewer agreement on abstract screening (Hanegraaf et al., 2024), so the ensemble is best read as a ranker rather than a binary classifier at a fixed threshold.

### Supplement J: Zero-label ranking versus ASReview active learning

We compared the universal zero-label ensemble ranking with `ASReview` (van de Schoot et al., 2021), a widely used active-learning system for study screening. ASReview was run with its default configuration (`elas_u4`: TF-IDF features, a support-vector classifier, certainty-based querying, and balanced training) on the same deduplicated corpora. Each run was initialised with one randomly selected relevant and one randomly selected irrelevant record, representing the minimal pilot typically required for active learning, and each review was repeated over ten random pilot selections. Both approaches were evaluated using the same recall definition used throughout this study, allowing direct comparison of the workload required to reach a specified recall target. Table S10 reports the workload required to achieve 95% recall.

The universal ranking achieved lower workload than ASReview on seven of the ten reviews and a lower mean workload overall (39% versus 42%). ASReview required less workload on *Coral Restoration*, *Coastal Wetlands Megafauna*, and *Habitat Restoration*, with gains of up to fourteen percentage points. In contrast, the largest differences favouring the universal ranking were observed for *Noise Pollution* (17 percentage points) and *Light Pollution* (14 percentage points). As ASReview updates its ranking as additional labels become available, these results reflect performance under a minimal pilot and the default model configuration rather than a tuned active-learning workflow. The comparison indicates that zero-label universal ranking can achieve screening efficiency comparable to supervised active learning under a minimal-pilot setting, while eliminating the need for review-specific training labels.

| **Review** | **Prev.** | **LLM** ($\to$ 95%) | **ASReview** ($\to$ 95%) | $\Delta$ **(LLM−ASR)** |
|---|---|---|---|---|
| Artificial | 0.02 | **0.16** | 0.30 [0.30, 0.30] | −0.14 |
| Habitat Restoration | 0.08 | 0.50 | **0.37** [0.37, 0.37] | +0.14 |
| CBFM | 0.09 | **0.38** | 0.40 [0.39, 0.41] | −0.02 |
| Coral Restoration | 0.10 | 0.31 | **0.27** [0.27, 0.27] | +0.04 |
| Light Pollution | 0.11 | **0.45** | 0.58 [0.54, 0.62] | −0.14 |
| Spatial Context | 0.12 | **0.36** | 0.38 [0.38, 0.38] | −0.02 |
| Coastal Wetlands Megafauna | 0.13 | 0.43 | **0.36** [0.36, 0.37] | +0.06 |
| Habitat Effect | 0.14 | **0.58** | 0.62 [0.62, 0.63] | −0.04 |
| Invasive | 0.15 | **0.41** | 0.43 [0.43, 0.44] | −0.03 |
| Noise Pollution | 0.25 | **0.29** | 0.46 [0.42, 0.48] | −0.17 |
| **Mean** | | **0.39** | 0.42 | |

Table S10: Workload to reach 95% recall (fraction of the corpus a human screens) for the universal zero-label LLM ranking versus ASReview active learning, per review, sorted by accept prevalence. ASReview values are the mean over ten random pilots with the 10th–90th percentile spread in brackets. Lower is better; the lower of the two is shown in bold. The LLM ranking is the lower-workload option on 7 of the 10 reviews and lower on average, despite using no labels, whereas ASReview learns from every label revealed during screening.

### Supplement K: Score-thresholding analysis

This supplement backs up our decision to use rank-position stopping rules instead of a fixed score threshold. Throughout we use the universal ranking from RQ2, the four-LLM *mean* ensemble {gemma3, gpt-oss, mistral-small3, qwen3} averaged over all three replicates per LLM. Every analysis uses the same aggregated score $A$, and only the review changes.

**Per-review $95\%$-recall thresholds vary by a factor of three.** The lowest threshold $\tau_{95}$ that still recovers $\geq 95\%$ of a review's relevant records (definition in Supplement L) ranged from $32.3$ on *Coastal Wetlands Megafauna* to $87.5$ on *Noise Pollution* under the same ranking (Figure S5), so no single fixed cutoff transfers across reviews.

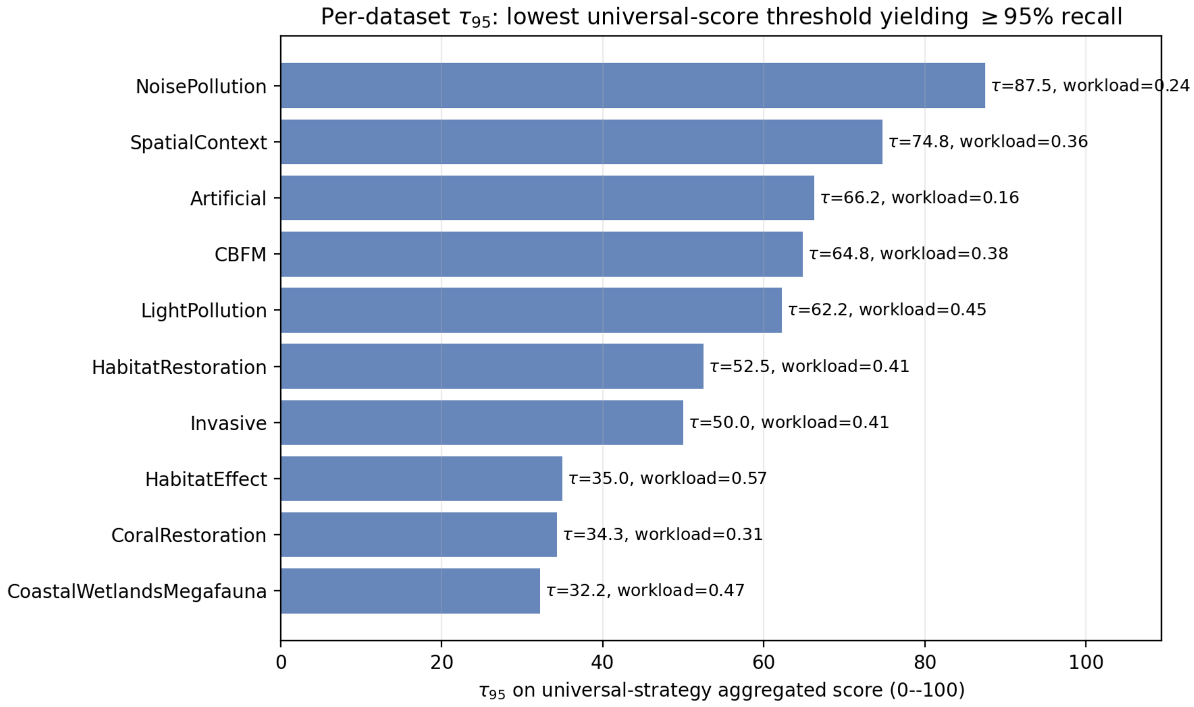


Figure S5: Per-review lowest universal-score threshold $\tau_{95}$ that achieves $\geq 95\%$ recall on the full review, sorted from highest to lowest. The same universal ranking strategy yields thresholds spanning $32$ to $88$ across the ten reviews, with implied workloads from $16\%$ (*Artificial*) to $57\%$ (*Habitat Effect*).

**Implication.** This is the point our rejection of score thresholding rests on. The threshold that reaches a given recall is review-specific even under a fixed ranking, so no single score cutoff transfers across reviews. The rank-position stopping rules in RQ4 avoid the problem. They are invariant to monotonic transformations of the score, so an uncalibrated score scale does not matter, and the audit-based ones (Bayesian, recall-confidence, order-statistics, Chao $r_r$) probe the unscreened tail during screening instead of fixing a single decision boundary up front.

### Supplement L: Formal mathematical definitions

The main text describes the stopping rules in plain language. This supplement gives the formal definitions for readers who want to reproduce or audit the rules.

| **Family** | **Rule** | **What it does** | **Settings tested** |
|---|---|---|---|
| Sequence-only | Consecutive negatives | Stop after a run of irrelevant records. | $L \in \{50, 100, 250, 500\}$ |
| | Percentage | Stop after a fixed fraction of the corpus has been screened. | $f \in \{10\%, 20\%, 25\%, 50\%\}$ |
| Periodic-audit | Bayesian | Audit the unscreened tail every 50 records and stop when the posterior says recall is very likely above target. | — |
| | Recall-confidence | Frequentist counterpart of Bayesian. | — |
| | Order-statistics | Stop when even a worst-case bound on remaining positives sits within the recall budget. | — |
| | Chao $r_r$ | Use the Chao1 population-size estimator on audit data; stop when found positives clearly outpace estimated remaining. | — |
| Combined heuristic | SAFE | Require a minimum coverage, a run of irrelevant records, and enough positives found to all hold at once. | coverage and run-length grids above; spot-check multiplier fixed at 2 |

Table S11: The seven stopping rules evaluated in the main text, plus the pre-calculated ideal-stopping-point rule used as a workload lower limit. Parameter grids for the sequence-only rules and SAFE are reported here; audit-based rules have no swept parameters (audit cadence and target recall are fixed at 50 records and 95% respectively). Formal mathematical definitions for the audit-based rules are given below. The model-aware rules, which did not improve on SAFE, are described in Supplement Q.

**Notation used in this supplement.**

- $N$: total number of labelled records in a review.
- $N_+$: true number of relevant (positive) records in the review.
- $\widehat{N}_+$: a reviewer's estimate of $N_+$, e.g. from a pre-screen spot-check.
- $j$: position in the ranked list, 1-indexed (so $j = 1$ is the highest-scored record).
- $f_j$: number of relevant records found by position $j$.
- $U_j = N - j$: size of the unscreened tail at position $j$.
- $A$: aggregated relevance score per record (the output of a ranking strategy on the 0–100 scale).
- $r$: number of replicates per LLM used by an ensemble strategy ($r \in \{1, 2, 3\}$).

**Recall budget.** For target recall $R^\star = 0.95$, after $f_j$ relevant records have been found by position $j$, the largest number of as-yet-unseen relevant records that can still be missed without dropping overall recall below $R^\star$ is

$$r_0(j) \;=\; \left\lfloor \frac{(1-R^\star)\, f_j}{R^\star} \right\rfloor.$$

This is the recall budget the audit-based rules (Bayesian, recall-confidence, order-statistics, Chao $r_r$) compare against the estimated number of unseen positives in the unscreened tail.

**Pre-calculated stopping target (ideal stopping point).** The pre-calculated rule stops at the smallest position $j$ at which the cumulative number of relevant records found, $f_j$, satisfies

$$f_j \;\geq\; K \;=\; \lceil R^\star \cdot N_+ \rceil.$$

Because $N_+$ is unknown to a real reviewer, this rule is reported only as a labels-known workload lower limit, not as a competitor.

**SAFE.** SAFE stops at the smallest $j$ at which all three conditions hold simultaneously:

1. minimum-coverage criterion: $j \geq \lceil p_{\min} N \rceil$, with $p_{\min}$ chosen from the grid $\{0.05$, $0.10$, $0.20$, $0.25$, $0.50\}$;
2. consecutive-negatives gate: the last $L$ records were all negatives, with $L$ chosen from $\{50, 100, 250, 500\}$;
3. spot-check criterion: $f_j \geq r_r \cdot \widehat{N}_+^{\text{spot}}$, where $r_r = 2$ and $\widehat{N}_+^{\text{spot}} = \text{round}(\hat{p}\, N)$ is derived from a 200-record random pilot's prevalence $\hat{p}$.

**Bayesian audit rule.** Every 50 records, draw an audit sample of size $m$ from the unscreened tail with $y$ positives observed. Update a Beta–Binomial predictive distribution from a uniform prior:

$$\Pr\big(\text{tail residual} \leq r_0(j) \,\big|\, \text{audit data}\big)$$

is computed under a $\text{Beta}(1+y,\, 1+m-y)$ posterior on tail prevalence. Stop when this probability exceeds $1-\alpha$ with $\alpha = 0.05$.

**Recall-confidence rule (frequentist).** Same audit structure as the Bayesian rule. At each audit, let $y$ positives be observed in $m$ tail draws. Compute the one-sided upper $1-\alpha$ binomial confidence bound on tail prevalence $\hat{p}^{\text{upper}}$ and the implied upper bound on remaining positives $\hat{p}^{\text{upper}} U_j$. Stop when $\hat{p}^{\text{upper}} U_j \leq r_0(j)$ with $\alpha = 0.05$ (Callaghan and Müller-Hansen, 2020).

**Order-statistics rule.** Every 50 records, compute the upper 95% Clopper–Pearson confidence bound on the prevalence of positives in the unscreened tail, multiply by the tail size $U_j$, and stop when even that worst-case estimate is less than or equal to the recall budget $r_0(j)$.

**Chao $r_r$ rule.** Use the Chao1 estimator to estimate the unseen-positive count from audit data. Let $f_1, f_2$ be the singleton and doubleton counts of positives in the audit samples drawn from the tail. The Chao1 lower bound on tail positives is $\widehat{N}_+^{\text{tail}} = y + f_1^2/(2f_2)$ (with the standard small-$f_2$ adjustment, $\widehat{N}_+^{\text{tail}} = y + f_1(f_1 - 1)/2$ when $f_2 = 0$). Let the Chao1-implied total positives be $\widehat{N}_+ = f_j + \widehat{N}_+^{\text{tail}}$. Stop when either (a) $f_j \geq r_r \cdot \widehat{N}_+$ with $r_r = 2$, or (b) the implied unseen-positive count $\widehat{N}_+ - f_j$ is within the recall budget $r_0(j)$.

**GLM (model-aware) rules.** Let $A_i \in [0, 100]$ be the universal-strategy aggregated score for record $i$. Begin with a $100$-record calibration sample $S_0$ formed by drawing $10$ records from each of the ten quantile bins (deciles) of the score distribution. Walk the unscreened pool $U$ in descending order of $A$. After each labelled record, refit a logistic regression

$$\log \frac{\Pr(y = 1 \mid A)}{1 - \Pr(y = 1 \mid A)} = \beta_0 + \beta_1 A$$

on the cumulative labelled set, and compute the expected positives remaining in the unscreened pool as $\hat{E}[\text{remaining}] = \sum_{j \in U} \hat{p}_j$ where $\hat{p}_j$ is the GLM's predicted probability for record $j$. The three GLM variants differ only in the stopping condition: **GLM recall** stops when $f/(f + \hat{E}[\text{remaining}]) \geq R^\star$ (here $f$ is the cumulative count of positives found, including the calibration sample); **GLM expected-count** stops when $\hat{E}[\text{remaining}] < K$ for $K \in \{1, 2, 5, 10\}$; **GLM proportion** ignores the GLM and stops when $f \geq p \cdot N$ for $p \in \{0.05, 0.10, 0.20\}$.

**Per-review $95\%$-recall threshold ($\tau_{95}$).** Used in Supplement K. On the full review, $\tau_{95}$ is the lowest aggregated-score cutoff $\tau$ such that the fraction of the review's positives with $A \geq \tau$ is at least $0.95$.

### Supplement M: Model provenance and inference environment

All results are conditional on the five open-weight models and quantisation levels listed in Table S12, served with Ollama on an NVIDIA H100 GPU. For reproducibility, we report the content digest of each model's Ollama manifest layer (`registry.ollama.ai/libra` which uniquely identifies the corresponding quantised weight blob. The `latest` tags used for *gpt-oss* and *mistral-small3.2* resolved to the 20B and 24B variants, respectively (*gpt-oss* records `from: gpt-oss:20b` in its manifest). Complete 64-character digests are provided with the released code.

| Model (tag served) | Params | Quant. | License | Model-layer digest |
|---|---|---|---|---|
| deepseek-r1:14b | 14B (reasoning) | Q4_K_M | MIT† | `sha256:6e9f90f0...` |
| gemma3:27b | 27B | Q4_K_M | Gemma Terms | `sha256:e796792e...` |
| gpt-oss:20b | 20B (MoE) | MXFP4 | Apache-2.0 | `sha256:b112e727...` |
| mistral-small3.2:24b | 24B | Q4_K_M | Apache-2.0 | `sha256:41a5b0c3...` |
| qwen3:30b-a3b | 30B (3B act., MoE) | Q4_K_M | Apache-2.0 | `sha256:78b329e7...` |

Table S12: Open-weight models used in this study, including quantisation, licence, and abbreviated model-layer digest. Results are conditional on the specified model versions and inference precisions. Full Ollama digests and software versions are provided in the released reproducibility materials (`environment.md`).

### Supplement N: Screening prompts

#### Initial Screening Prompt

You are screening a list of academic records based on the following inclusion criteria:

1. Must focus on fish (bony or cartilaginous species).
2. Must describe a study conducted in a marine or coastal environment (including estuaries, bays, lagoons, or seagrass beds).
3. Must include a comparison between two or more habitat types, habitat conditions, or habitat characteristics (e.g., natural vs artificial, restored vs degraded, native vs invasive, complex vs simple).
4. Must include measurements of fish fitness using behavioural or physiological indicators (e.g., feeding rate, boldness, growth, cortisol, survival, GSI, immune function, etc.).

Please make a binary decision for each record: "include" or "not_include". Provide a brief explanation for your decision.
Respond ONLY with a JSON array of objects in the following format:

```
[
  {
    "id": "paper_0",
    "decision": "include",
    "explanation": "Explanation here"
  },
  ...
]
```

#### Standard Screening Prompt (dynamic template)

The prompt is generated dynamically per review. Placeholders in angle brackets are filled at screening time: «`SCOPE_SUMMARY`» is the review's one-line scope; «`INCLUSION_CRITERIA`» expands to the review's numbered inclusion criteria; «`CLARIFICATIONS`» is the optional habitat-clarification block (used for *Artificial*, *Habitat Effect*, *Habitat Restoration* only); $n$ is the number of inclusion criteria for the review (3, 4, or 5 across our reviews); $W = \mathrm{round}(100/n)$ is the per-criterion maximum (33, 25, or 20 points); and the per-criterion scale thresholds $p_{20}, p_{40}, p_{80}$ scale with $W$.

```
You are screening a list of academic records.
Scope summary: <<SCOPE_SUMMARY>>

Apply the following inclusion criteria to the Title & Abstract.
Score **each criterion separately**; then sum to get the 0-100
  relevance score:
<<INCLUSION_CRITERIA>>

Clarifications:                                  (optional block)
<<CLARIFICATIONS>>

Scoring guidance (dynamic):
- There are <n> criteria. Each criterion is worth ~<W> points (100
  total).
- Award partial credit when a record partially meets a criterion or is
  ambiguous but suggestive.
- Per-criterion scale (proportional to the per-criterion maximum):
  - 0 points: Criterion clearly not met.
  - ~<p20>-<p40> points: Weakly supported or only tangentially
    addressed.
  - ~<p40>-<p80> points: Moderately supported but lacking clarity or
    completeness.
  - ~<p80>-<W> points: Clearly and fully met.

Return a final **relevance score (0-100)** equal to the sum of per-
  criterion
points. Provide a brief explanation referencing which numbered
  criteria are
satisfied, uncertain, or not met.

Respond ONLY with a JSON array of objects in the following format
(do not repeat title/abstract text):
[
  {
    "id": "paper_0",
    "explanation": "Brief rationale referencing each criterion and its
                   partial score (e.g., C1: 18/25, C2: 12/25, C3:
                     0/25,
                   C4: 22/25 - totals sum to relevance score)",
    "relevance": "50"
  },
  ...
]

Ensure to include ALL paper_id values you were given.
```

For a review with $n = 4$ criteria, the rendered weights are $W = 25$ and per-criterion scale boundaries $(p_{20}, p_{40}, p_{80}) = (5, 10, 20)$. The implementation is in `scripts/utils/helper.py` (function constructing `base_prompt`).

### Supplement O: The benchmark as a candidate resource for ecological screening

The evaluation in this paper rests on a collection of ten fully human-labelled title-and-abstract screening corpora, together $19{,}155$ labelled records (Table 1 in the main text). Six correspond to published systematic reviews and four are in preparation. We document the collection here and set out why we believe it could serve as a shared benchmark for LLM-assisted screening in ecology and environmental science, alongside the two external terrestrial benchmarks we also test against ((Nykvist et al., 2025; Nawrath et al., 2026)).

**Composition and documentation.** Each record is associated with a human include/exclude decision made at the title-and-abstract screening stage, the task targeted by the ranking methods evaluated here. The full inclusion criteria for all reviews are reproduced in Supplement P, and the released dataset includes all per-record, per-model, and per-replicate LLM relevance scores together with the corresponding model-manifest digests (Supplement M). These resources permit direct evaluation of alternative ranking, aggregation, and stopping methods using the same inputs and labels analysed in this study. One review (*Habitat Effect*) includes an expert re-review of records with the largest ensemble–screener disagreements, and three reviews (*Light Pollution*, *Spatial Context*, and *Coral Restoration*) include stratified blinded re-audits by two independent readers, providing empirical estimates of label uncertainty (Supplement D).

**Benchmark potential.** Public screening benchmarks are abundant in biomedicine and information retrieval but comparatively scarce in ecology and environmental science. This collection contributes a domain-specific benchmark for title-and-abstract screening in environmental evidence synthesis. The included reviews span a range of ecological topics and screening criteria, providing realistic variation in scope, terminology, and relevance definitions. Each corpus is paired with original reviewer decisions and eligibility criteria, allowing evaluation of the complete rank-and-stop screening task rather than a proxy classification objective. The reviews also vary substantially in size and prevalence, reducing dependence on any single operating point. Finally, the release of the underlying LLM score matrices enables reproducible comparison of alternative aggregation and stopping methods without requiring model re-execution.

**Availability.** The six published reviews are citable through their source papers (Table 1), and an example corpus (*CBFM*) ships with the `screenllm` package as a runnable vignette. The remaining in-preparation reviews are released with their parent publications and are available from the authors on request in the interim, anonymised where the underlying review is not yet public.

### Supplement P: Per-review inclusion criteria

This supplement reproduces the inclusion criteria used for each of the ten labelled reviews in human-readable form. Each review's inclusion criteria are the rules the human review leads supplied; the LLM ensemble screens against the same numbered list inserted into the standard prompt template (Prompts section). The habitat clarifications block at the end is shared across the three habitat-focused reviews (*Artificial*, *Habitat Effect*, *Habitat Restoration*).

##### *Artificial*

**Scope:** Articles that explore individual fitness or morphometric measure response to marine habitats. Comparisons can include habitat type (artificial vs. natural; seagrass vs. kelp) or habitat state (complex vs. simple; restored vs. degraded).

**Inclusion criteria:**

1. The Title and Abstract summarise a study that focuses on fish and/or sharks.
2. The study is conducted in a marine or coastal environment.
3. The study describes a comparison between two or more habitat types, habitat conditions, or habitat characteristics.
4. The study describes direct or indirect measurements of fish fitness using behavioural and physiological indicators or morphological traits.

##### *CBFM*

**Scope:** Articles potentially relevant to community-based fisheries management (CBFM) in Pacific Island contexts.

**Inclusion criteria:**

1. It is possible that the study includes a case study from one or more of: Cook Islands, Federated States of Micronesia, Fiji, Kiribati, Marshall Islands, Nauru, Niue, Palau, Papua New Guinea, Samoa, Solomon Islands, Tonga, Tuvalu, or Vanuatu.
2. It is possible that the study discusses fisheries and/or marine resource management.
3. It is possible that the study discusses a community-based approach.

##### *Coastal Wetlands Megafauna*

**Scope:** Articles that explore how marine megafauna associate with vegetated coastal wetland habitats, reporting empirical evidence of implicit or explicit habitat associations.

**Inclusion criteria:**

1. The study focuses on one or more marine megafauna taxa (dugongs, manatees, sea turtles, otters, seals, minks, crocodiles, alligators, sharks, rays, dolphins, porpoises).
2. The study includes one or more vegetated coastal wetland habitats (mangroves, seagrass, saltmarsh / tidal marsh).
3. The study reports empirical data indicating an implicit or explicit habitat association between megafauna and these habitats.

##### *Coral Restoration*

**Scope:** Articles that examine the performance of field-based coral reef restoration projects using active restoration interventions and monitoring at least one performance metric.

**Inclusion criteria:**

1. Title and Abstract describe a study conducted at a field-based coral reef restoration site (i.e., a natural reef setting, not an in-situ or ex-situ coral nursery).
2. Title and Abstract describe a project with the explicit goal of restoring, rehabilitating, or improving a coral reef ecosystem.
3. Title and Abstract describe the use of an active coral reef restoration intervention, such as substrate addition, coral transplantation/outplanting, larval enhancement, or similar treatments.
4. Title and Abstract describe monitoring or assessment of at least one restoration performance metric (e.g., coral survival, growth, recruitment, cover, biodiversity response, ecosystem function).

##### *Habitat Effect*

**Scope:** Articles that explicitly examine how surrounding or adjacent habitats influence the ecological performance of artificial reefs as marine habitats.

**Inclusion criteria:**

1. The study is conducted in a marine or estuarine environment.
2. The study examines fish assemblages or fish performance associated with artificial reefs or artificial hard structures functioning as reefs (e.g., purpose-built reefs, shipwrecks, oil and gas platforms).
3. The study explicitly tests, compares, or models how surrounding or adjacent habitats (e.g., seagrass, mangroves, natural reefs, soft sediments, pelagic environments) influence artificial reef performance, using spatial contrasts, gradients, controls, or mechanistic reasoning.
4. The study reports fish responses that reflect artificial reef performance, such as fitness-related, demographic, behavioural, physiological, or population-level indicators, and interprets these responses in relation to surrounding habitat context.

##### *Habitat Restoration*

**Scope:** Articles that examined a response of organisms at any level to habitat restoration efforts.

**Inclusion criteria:**

1. The study is conducted in saltmarsh, mangrove, seagrass, macroalgae forest, coral reef, or oyster reef ecosystems.
2. The study focuses on restoration actions directed toward habitat-forming species (e.g., planting, translocations, deploying structures to explicitly attract habitat formers, exotic species removal, restoring tidal flow to allow natural processes to reinitiate with respect to vegetation).
3. The study reports measured responses of non-habitat-forming animals that reflect population-level dynamics or individual-level performance.

##### *Invasive*

**Scope:** Articles that compared invasive species occupancy of artificial and natural habitats.

**Inclusion criteria:**

1. The study is conducted in a marine or estuarine environment (coastal waters, estuaries, bays, nearshore marine habitats; excludes freshwater).
2. The study examines invasive species.
3. The study includes both artificial structures (e.g., pilings, seawalls, pontoons, breakwaters, artificial reefs, or other built environments) and natural reefs (e.g., coral or rocky reefs). If natural reefs are not explicitly mentioned, accept if control sites likely represent natural reefs.
4. The study compares invasive species on actual artificial structures and natural reefs (or control sites).

##### *Light Pollution*

**Scope:** Articles examining the effects of light pollution on fitness of fish and invertebrates in the marine environment.

**Inclusion criteria:**

1. The study is conducted in a marine or coastal environment.
2. The study focuses on fish and/or sharks and/or marine invertebrates.
3. The study describes exposure to artificial light or light pollution.
4. The study describes direct or indirect measurements of fitness or individual performance using behavioural and/or physiological indicators, or morphological traits.

##### *Noise Pollution*

**Scope:** Articles examining the effects of marine noise pollution on community response, fitness, or individual performance of fish and invertebrates, restricted to empirical studies.

**Inclusion criteria:**

1. The study is conducted in a marine or coastal environment.
2. The study focuses on fish and/or sharks and/or marine invertebrates.
3. The study presents empirical data (excludes reviews and modelling studies).
4. The study describes exposure to marine noise pollution.
5. The study describes direct or indirect measurements of community response, or fitness or individual performance, using behavioural and/or physiological indicators, or morphological traits.

##### *Spatial Context*

**Scope:** Articles that examine aspects of seascape/habitat spatial configuration (such as patch size, distance between patches, and isolation) and how these relate to the fitness or performance of marine fish or invertebrates in submerged coastal habitats.

**Inclusion criteria:**

1. The study focuses on one or more habitat-dependent marine animals (fish or invertebrates).
2. The study was conducted in submerged coastal or marine habitats (e.g., reefs, seagrass, kelp, mangroves below tideline).
3. The study describes a comparison or contrast in seascape/spatial context between two or more sites/habitats (e.g., patch size, spacing, isolation).
4. The study reports animal responses to spatial-context variables, including fitness-related, demographic, physiological, population-level indicators, and/or habitat selection/use.

##### Habitat clarifications block (used by *Artificial*, *Habitat Effect*, *Habitat Restoration*)

For the purposes of these reviews:

- *Habitat* is a three-dimensional physical structure (biogenic or abiotic, natural or human-made) that provides space, surface, shelter, or food for biological communities.
- *Habitat conditions* refer to the physical state or integrity of the habitat (e.g., structural complexity, degradation, fragmentation), not environmental parameters such as water quality, pollution, temperature, or salinity.
- *Habitat characteristics* refer to inherent physical features of the habitat (e.g., substrate type, rugosity, vertical relief), excluding chemical or biological environmental factors.

#### Supplement Q: Model-aware stopping rules

Beyond the seven rules compared in the main text, we tested three stopping rules that use the LLM relevance score directly. Each starts with a 100-record decile-stratified calibration sample, then walks down the universal ranking and, after every labelled record, refits a logistic regression of relevance on the LLM score. The fitted model gives an expected number of relevant records still left in the unscreened pool (formal definitions in Supplement L). The three variants use that estimate differently. *GLM expected-count* stops when the expected remaining count drops below a threshold $K$, *GLM proportion* stops when the positives found reach a fixed fraction of the corpus, and *GLM recall* stops when the implied expected recall reaches the target. None improved on SAFE. At their tuned settings, GLM expected-count reached the $95\%$ target on $8/10$ reviews at $56\%$ mean workload, GLM proportion reached $9/10$ but read $89\%$ of the corpus on average, and GLM recall reached only $3/10$ because it stops too early. The rest of this supplement examines that last failure, which is the most informative about the limits of using the LLM score as a recall estimator.

The GLM-recall rule stops when the implied expected recall, $f/(f + \hat{E}[\text{remaining}])$, reaches a chosen target. Its workload therefore depends on whether the regression's plug-in estimate of recall at the stopping point is well calibrated. In the main Results we report the rule at a single calibration seed at target $0.95$, where it reached the target on only $3/10$ reviews. This supplement examines whether that under-stopping is a property of the rule or an artefact of using a single calibration draw.

We swept the recall target over $\{0.80, 0.85, 0.90, 0.95, 0.99\}$ and repeated each with $20$ independent decile-stratified calibration samples ($10$ per decile, $n = 100$ total). For each review, target, and seed we recorded both the recall the rule *predicted* at its stopping point and the recall it actually achieved on the full review. Figure S6 plots actual against predicted, faceted by review and coloured by target. The diagonal marks perfect calibration.

The pattern is mixed and review-dependent. The rule is well calibrated on *CBFM*, *Invasive*, *Light Pollution*, and *Noise Pollution*, where predicted and actual recall track each other across the target range with error bars covering the diagonal. On *Coral Restoration*, *Habitat Restoration*, and *Spatial Context* it is consistently *optimistic*, stopping where its predicted recall is at or near target but the actual recall is $0.07$–$0.15$ lower. *Habitat Effect* shows large error bars throughout. *Coastal Wetlands Megafauna* switches from optimistic at low targets to slightly conservative at $0.95$, indicating an unstable jump in the stopping position as the target tightens. *Artificial* is discontinuous, because its very low positive count ($N_+ = 73$ out of 3,846) leads the rule to threshold either at the first calibration positive or at the end of the ranking, with no intermediate behaviour.

**Interpretation.** The under-target outcome reported in the main text is therefore not an artefact of a single calibration seed. At target $0.95$, the median actual recall across $20$ seeds stays below $0.95$ on the four optimistic reviews, and on *Habitat Restoration* and *Spatial Context* it sits at $0.85$–$0.86$. The bias appears to depend on how the relationship between LLM score and relevance probability is structured in each review's tail. A $100$-record calibration sample is sufficient to fit the regression, but not, on the harder reviews, to characterise the deep-tail probability well enough for the plug-in recall estimate to be unbiased. The companion *GLM expected-count* rule stops on the

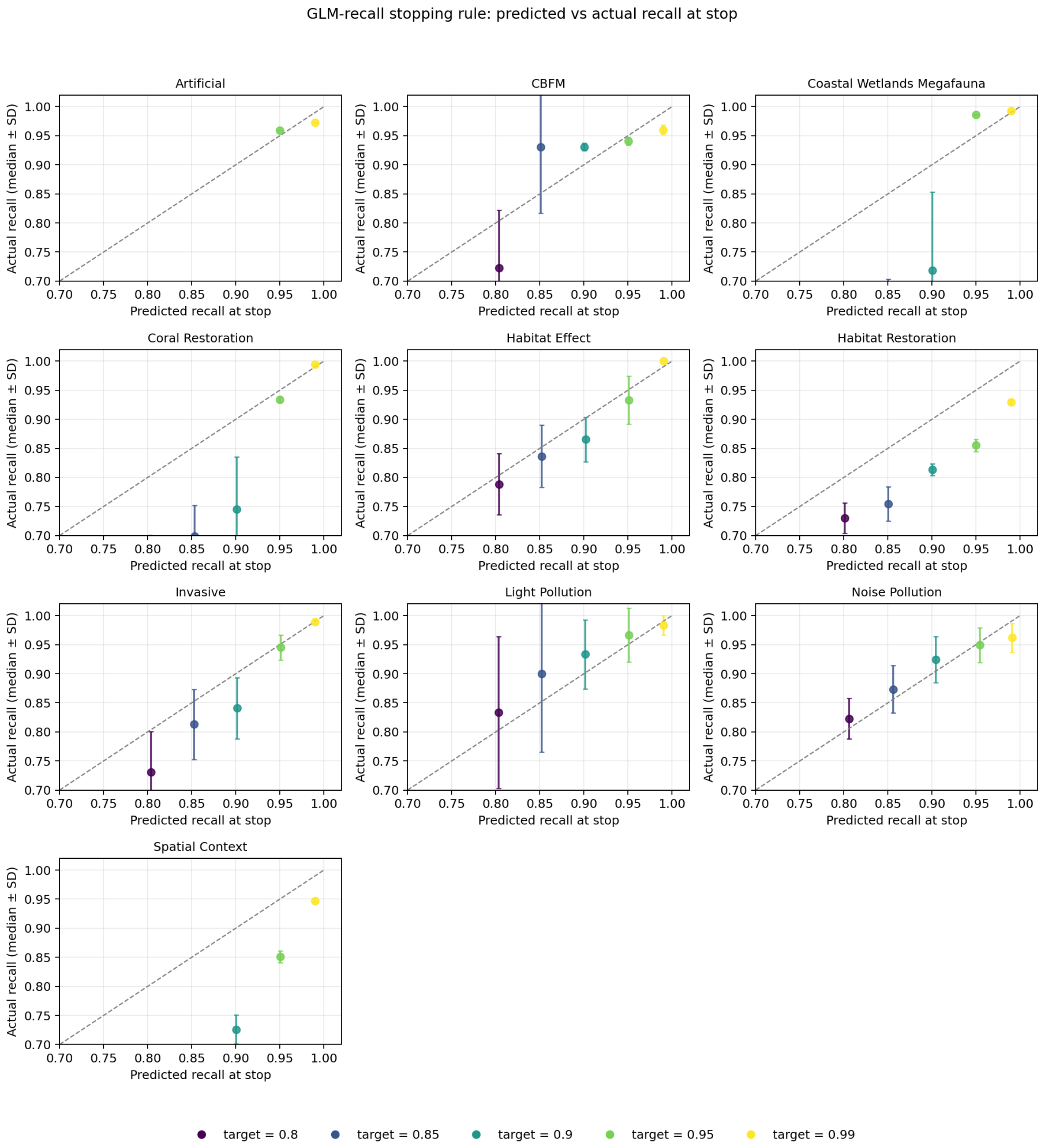


Figure S6: GLM-recall stopping rule, predicted vs. actual recall at the stopping point, per review. Each point is the median across 20 independent decile-stratified calibration draws at a fixed recall target ($n_{\text{per decile}} = 10$, $n_{\text{cal}} = 100$), and error bars are $\pm 1$ SD of actual recall across the 20 draws. The dashed line is perfect calibration. Points below it mean the rule is optimistic (predicted recall higher than actual), and points above it mean the rule is conservative.

unweighted count $\hat{E}[\text{remaining}] < K$, not the ratio, and is much less affected ($8/10$ at $K \in \{1, 2\}$). The ratio structure of the recall estimate amplifies tail-probability bias in a way the count does not.
